# Multi-task machine learning reveals the functional neuroanatomy fingerprint of mental processing

**DOI:** 10.1101/2023.11.30.569385

**Authors:** Zifan Wang, Yuzhong Chen, Jiadong Yan, Yi Pan, Wei Mao, Zhenxiang Xiao, Guannan Cao, Paule-J Toussaint, Weitong Guo, Boyu Zhao, Hailin Sun, Tuo Zhang, Alan C Evans, Xi Jiang

## Abstract

Mental processing delineates the functions of human mind encompassing a wide range of motor, sensory, emotional, and cognitive processes, each of which is underlain by the neuroanatomical substrates. Identifying accurate representation of functional neuroanatomy substrates of mental processing could inform understanding of its neural mechanism. The challenge is that it is unclear whether a specific mental process possesses a ‘functional neuroanatomy fingerprint’, i.e., a unique and reliable pattern of functional neuroanatomy that underlies the mental process. To address this question, we utilized a multi-task deep learning model to disentangle the functional neuroanatomy fingerprint of seven different and representative mental processes including Emotion, Gambling, Language, Motor, Relational, Social, and Working Memory. Results based on the functional magnetic resonance imaging data of two independent cohorts of 1235 subjects from the US and China consistently show that each of the seven mental processes possessed a functional neuroanatomy fingerprint, which is represented by a unique set of functional activity weights of whole-brain regions characterizing the degree of each region involved in the mental process. The functional neuroanatomy fingerprint of a specific mental process exhibits high discrimination ability (93% classification accuracy and AUC of 0.99) with those of the other mental processes, and is robust across different datasets and using different brain atlases. This study provides a solid functional neuroanatomy foundation for investigating the neural mechanism of mental processing.

**One-Sentence Summary:** There exists a functional neuroanatomy fingerprint to underlie the mental process.

## Introduction

The brain receives, encodes, stores, retrieves, and manipulates information which constitutes mental processing and delineates brain functions. Over two centuries ago, it was proposed that human-specific mental processing could be determined based on the shape of the skull (*1*). Although the theory of phrenology was discredited in scientific research, the hypothesis that different neuroanatomical regions are associated with distinct brain functions has been supported in a series of studies conducted thereafter (*2-6*). Consequently, brain atlases were established to understand the relationship between brain anatomy and function (*7,8*). A specific mental process is accomplished by multiple brain regions adhering to two basic organizing principles: integration and segregation (*9*). The regionally separated brain regions defined by brain atlases are functionally linked as brain networks to achieve specific mental process (*10-16*), while a single brain region may be involved in multiple brain networks corresponding to different mental processes (*17-20*). Interestingly, while there are cognitive and behavioral differences among individuals with different cultural and ethnic background (*21-23*), there is certain consistency of neural activation patterns under the same mental process across different populations (*24,25*). However, whether a specific mental process possesses a unique and reliable pattern of functional neuroanatomy which differentiates with other mental processes, is largely underexplored.

Since the phenomenon of high functional coherence between the left and right brain motor regions was discovered using functional magnetic resonance imaging (fMRI)-derived BOLD signals (*26*), fMRI has been widely adopted and combined with various data modeling approaches to characterize the mental processes and the associate neuroanatomical regions. For example, General Linear Model (GLM) is commonly used to identify activated brain regions under specific task paradigms (*27*). Matrix decomposition methods such as Independent Component Analysis (ICA) (*28*), Principal Component Analysis (PCA) (*29*), and Dictionary Learning (DL) (*30*) are adopted to decompose the spatiotemporal patterns of brain activities. Given the powerful ability of deep learning models in extracting high-dimensional nonlinear data features, recent studies have attempted to use advanced deep learning models to characterize fMRI signals and associated neuroanatomical patterns under specific mental process (*31*).

Among various measures of neural activities, functional connectivity (FC) is a crucial measure to characterize the functional coherence between neuroanatomical regions (*32*) during a mental process. The widely known concept of “functional brain fingerprint” (*33*) refers to the unique and reliable patterns of whole-brain FC which can effectively identify individuals using brain functional imaging techniques such as fMRI. The effectiveness and necessity of adopting whole-brain FC pattern to represent mental processes is also justified in previous studies reporting that even minor changes in task paradigms can alter the functional activity of the whole brain (*34*).

In order to explore the unique and reliable pattern of functional neuroanatomy for a mental process, and inspired by the concept of “functional brain fingerprint”, in this study, we adopted an advanced multi-task deep learning classification model to disentangle the functional neuroanatomy fingerprint of various mental processes based on task-based fMRI data. Specifically, taking the whole-brain FC patterns of various mental processes in a large cohort of subjects as model inputs, the Vision Transformer (ViT)-based (*35*) multi-task model aims to identify the unique functional neuroanatomy pattern (*36*) for each mental process with high discrimination accuracy with the others. Using the publicly available two independent datasets Human Connectome Project (HCP) and Chinese Human Connectome Project (CHCP) of 1235 subjects with seven task-based fMRI data representing seven different mental processes (Emotion, Gambling, Language, Motor, Relational, Social, and Working Memory), we hypothesize that each of the seven mental process possesses a ‘functional neuroanatomy fingerprint’ consistently across different individuals of two independent datasets. We also hypothesize that the identified functional neuroanatomy fingerprint is represented by a unique set of functional activity weights of whole-brain regions characterizing the degree of each region involved in a specific mental process, and exhibits well discrimination ability with those of other mental processes.

## Results

### Functional Neuroanatomy Fingerprint of Mental Process

We first identify the functional neuroanatomy fingerprint for each of the seven mental processes (Emotion, Gambling, Language, Motor, Relational, Social, and Working Memory) using the Human Connectome Project S1200 young dataset (HCP-YA) of 1007 subjects. To ensure the stability of model performance (Fig. 1A), we train and test the seven-class classification model using five-fold cross-validation with six different random seeds, resulting in 5×6=30 trained models in total corresponding to thirty fingerprints for each mental process (Fig. 1B). The functional neuroanatomy fingerprint achieves 93% averaged classification accuracy and area under the curve (AUC) of 0.99 among the seven mental processes (Fig. 1C), and exhibits stable pattern under different types of classification models (Supplemental Fig. 1), demonstrating the uniqueness and reliability of the fingerprint for each mental process.

**Fig. 1.**
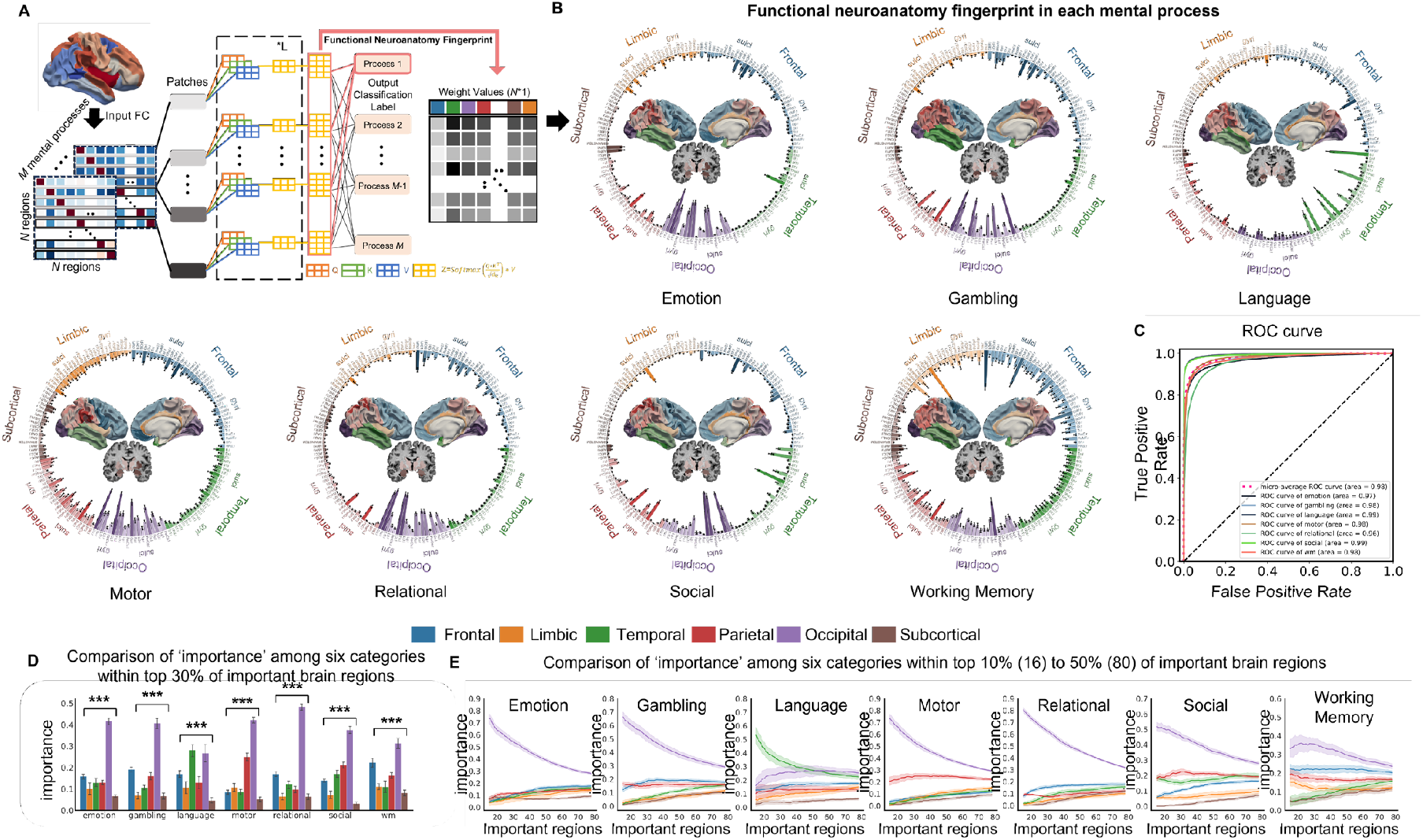
Functional neuroanatomy fingerprint for each of the seven mental processes. **(A)** Identification of functional neuroanatomy fingerprint based on the proposed multi-task deep learning model. The model inputs are the whole-brain functional connectivity matrices of the seven mental processes in all subjects, and the output is the seven-class classification label as well as the associated fingerprint contributing to the classification. **(B)** Representation of the functional neuroanatomy fingerprint in each mental process as a set of functional activity weight values of all neuroanatomical regions defined by the Desikan-Killiany (DK) brain atlas. Note that each value is represented as mean±std based on the thirty trained models. **(C)** ROC curves for each of the seven mental processes. **(D)** Comparison of ‘importance’ (defined as normalized ratio of involved important regions) among the six categories (five cortical lobes and one subcortical area) within top 30% of important brain regions in each mental process. **(E)** Comparisons of ‘importance’ among the six categories in each mental process under different thresholds of important brain regions from top 10% (16 brain regions) to top 50% (80 brain regions) with a 0.5% interval. The data is represented as the mean value with 95% confidence intervals, *** indicates p<0.001, and ns indicates not significant in six-way ANOVA. FDR correction is used for multiple comparisons.

The functional neuroanatomy fingerprint is represented as a set of functional activity weight values of each neuroanatomical region defined by the Desikan-Killiany (DK) brain atlas in Fig. 1B. More visualizations are provided in Fig. 2. We see that the identified functional neuroanatomy fingerprint for each mental process is stable in terms of low standard deviation of functional activity weight value in each region among the thirty trained models. We group the whole-brain neuroanatomical regions into six categories including five cortical lobes (frontal, parietal, temporal, occipital, and limbic) and one subcortical area for interpretation facilitation. We see that the identified fingerprint is closely related to the associated task paradigm in each mental process. For example, the identified fingerprint of Emotion exhibits large functional activity weights in amygdala which are crucial for emotional processing; the fingerprint of Language possesses large functional activity weights in temporal lobe which is responsible for language processing. Based on the functional activity weight value in each region, we select top *k%* regions with largest weight values as the important regions contributing to the fingerprint, and define the ‘importance’ value as normalized ratio of involved important regions in each of the six categories (five lobes and one subcortical area). Figs. 1D-1E demonstrate that the occipital lobe consistently exhibits significantly larger ‘importance’ than the other five categories (six-way ANOVA, p<0.001, FDR corrected, post-hoc t-test) under different thresholds of top *k%* important regions and across all seven mental processes. Since all of the seven task paradigms involve visual stimuli (e.g., images, videos, and text) or visual cues and feedbacks, it is reasonable that there is strong functional activity weight of visual regions (*37*) in the functional neuroanatomy fingerprint of each mental process.

**Fig. 2.**
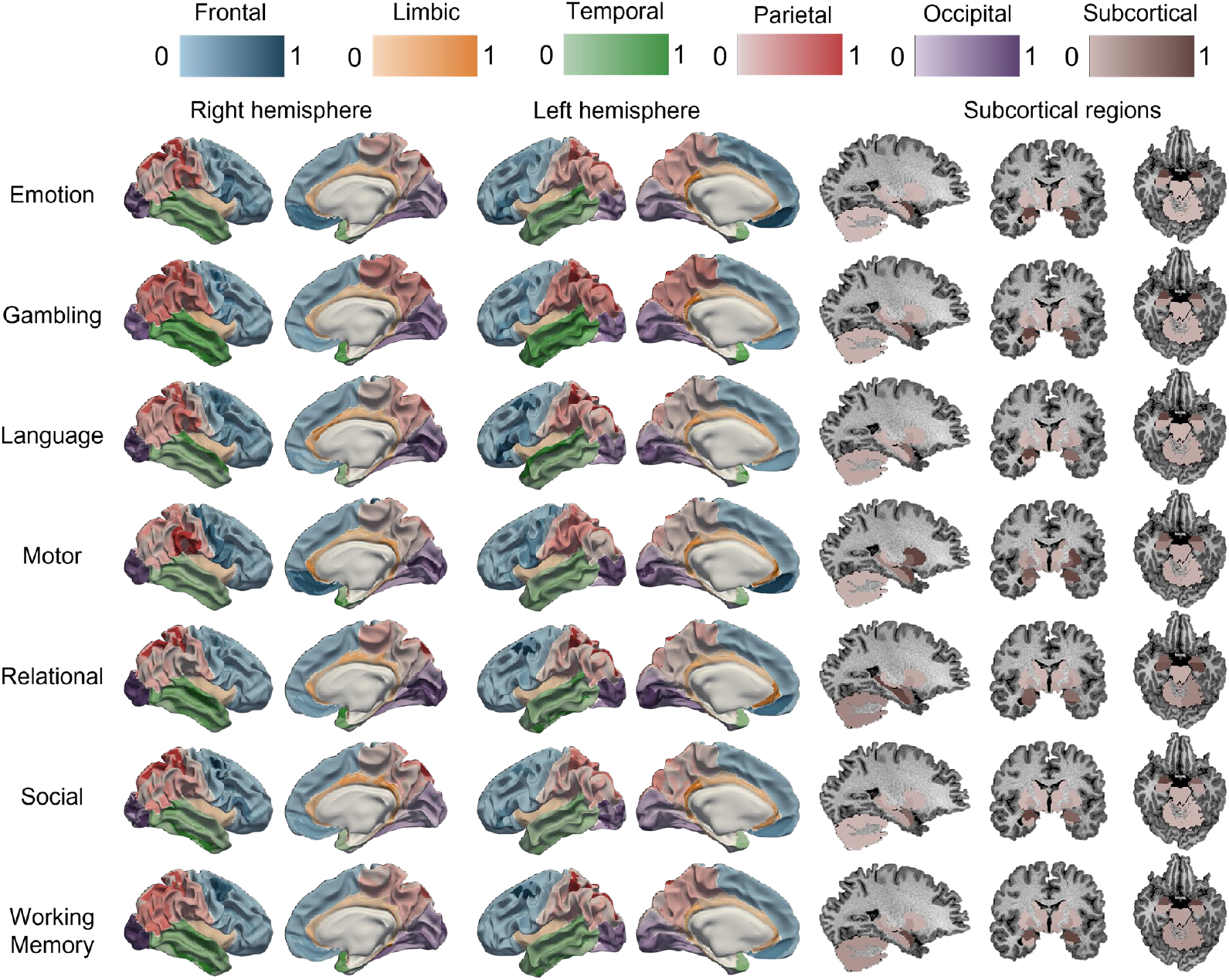
Distribution of the functional activity weight values in the six categories based on the HCP dataset and DK atlas. Each category (five lobes and one subcortical area) is represented by a unique color while larger/smaller weight values correspond to darker/lighter color.

### Commonality and Uniqueness of Functional Neuroanatomy Fingerprint among Different Mental Processes

We then investigate the commonality and uniqueness of identified functional neuroanatomy fingerprint among the seven mental processes. Fig. 3A highlights the common (exist in all seven mental processes) and unique (not in all seven ones) neuroanatomy regions in blue and red respectively. The common regions are mainly located in occipital and parietal lobes, with a few in the left frontal lobe and amygdala, while the unique regions are widely distributed across whole-brain. We further divide each cortical region into fine-scale gyri and sulci parts, and re-categorize each region into one of four categories based on whether its gyral/sulcal part is involved in top *k%* important regions with largest functional activity weight values in the fingerprint: 1. Gyri/sulci are both involved; 2. Gyri/sulci are both not involved; 3. Only gyri are involved; 4. Only sulci are involved. As shown in Fig. 3A, in occipital lobe, the gyri and sulci of lateral occipital (LO) region are both involved across all seven processes. However, in other regions of the occipital lobe such as pericalcarine, fusiform, and lingual regions, sulci are more involved than gyri. In parietal lobe, the sulci of the superior parietal region are involved in all seven mental processes, whereas the gyri of this region exclusively involve in Language, Relational, and Social ones. The sulci of the inferior parietal region are involved in all mental processes except for Motor, while the gyri are only involved in Language and Social. In temporal lobe, the region distribution is similar among all mental processes except for Language, where the superior temporal gyri and temporal pole are primarily involved. An intriguing difference is found in frontal lobe, particularly in the medial orbitofrontal region of the left hemisphere, where the sulci are consistently involved across all seven mental processes, while the gyri are not involved in Relational and Social. We also observe widespread involvement of the inferior frontal region across most mental processes. In limbic lobe, gyri are more involved than sulci in contrast to the other four cortical lobes. These findings highlight different involvement between gyri and sulci in mental processing, i.e., sulci play a more important role than gyri in discriminating different mental processes.

**Fig. 3.**
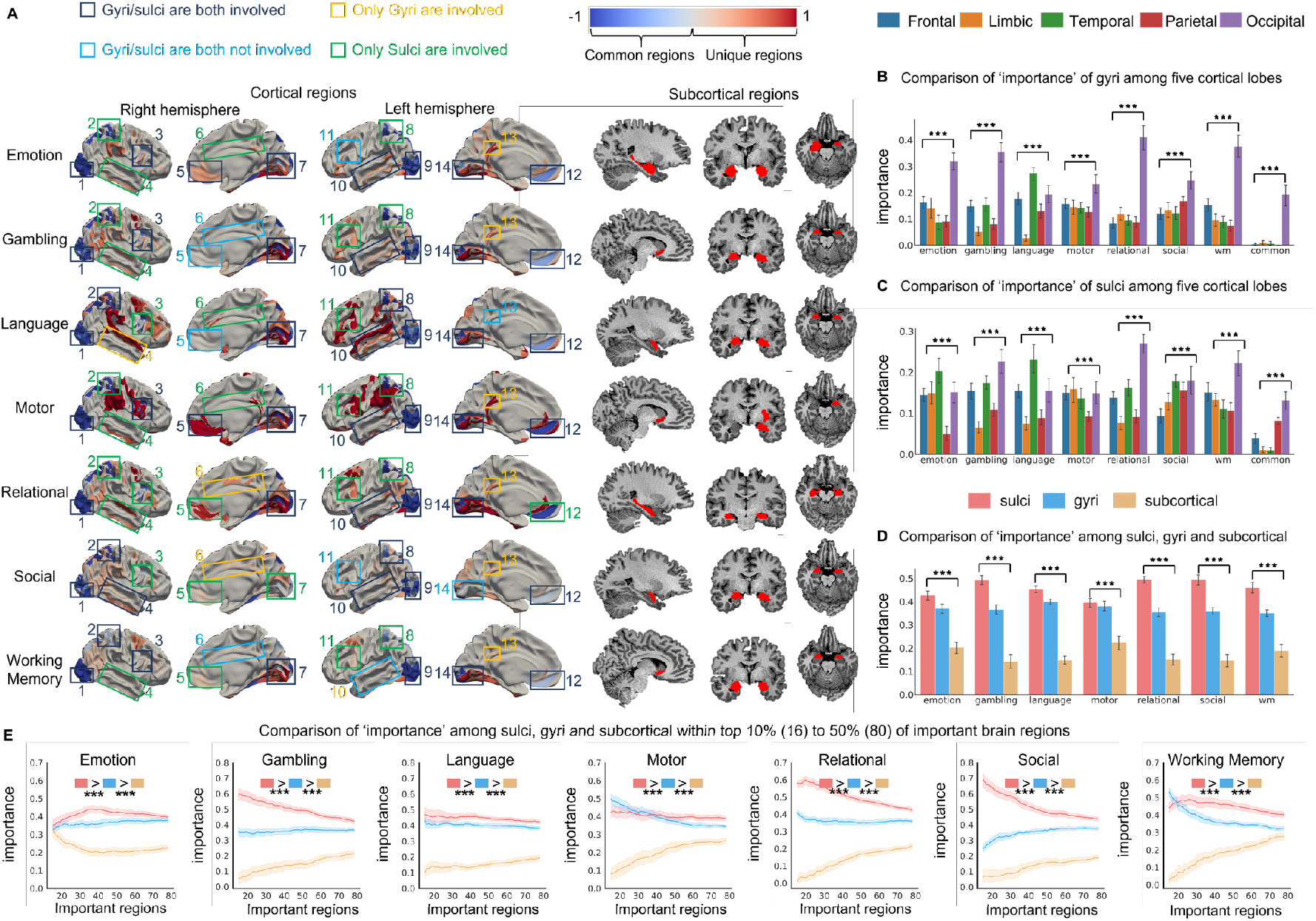
Commonality and uniqueness of involved neuroanatomy regions in fingerprint among different mental processes. **(A)** Distribution of the common (exist in all seven mental processes) and unique (not in all seven ones) neuroanatomy regions colored in blue and red respectively. Top *30%* important regions with largest functional activity weight values are used as example. Fourteen representative regions in each of the four categories based on the involvement of gyri/sulci are highlighted using four different colored boxes. **(B)** Comparison of ‘importance’ value of gyri among the five cortical lobes in the unique and common regions, respectively using top 30% of important brain regions. **(C)** Comparison of ‘importance’ value of sulci among the five cortical lobes in the unique and common regions, respectively using top 30% of important brain regions. **(D)** Comparison of ‘importance’ value among gyri, sulci, and subcortical area in each of the mental process using top 30% of important brain regions. **(E)** Comparison of ‘importance’ value among gyri, sulci, and subcortical area in each of the mental process under different thresholds of important brain regions from top 10% (16 brain regions) to top 50% (80 brain regions) with a 0.5% interval. The data is represented as the mean value with 95% confidence intervals, *** indicates p<0.001, and ns indicates not significant in multi-way ANOVA. FDR correction is used for multiple comparisons.

We further report significant difference (five-way ANOVA, p<0.001, FDR corrected, post-hoc t-test) of ‘importance’ value of gyri/sulci among the five cortical lobes in the unique and common regions, respectively. Taking top *30%* important regions with largest functional activity weight values as example, Fig. 3B shows that gyri of occipital lobe exhibit largest ‘importance’ in the unique regions of most mental processes except for Language, as well as in the common regions in Fig. 3A. Fig. 3C shows that sulci of occipital lobe exhibit largest ‘importance’ in the unique regions of Gambling, Relational, and Working Memory, as well as in the common regions, while sulci of temporal lobe have largest ‘importance’ in the unique regions of Emotion, Language, and Social. Figs. 3D-3E further show that sulci consistently exhibit significantly larger ‘importance’ than gyri and subcortical area (three-way ANOVA, p<0.001, FDR corrected, post-hoc t-test) under different thresholds of top *k%* important regions and across all seven mental processes.

### Robustness of Functional Neuroanatomy Fingerprint across Different Datasets and Brain Atlases

We assess the robustness of identified functional neuroanatomy fingerprint for each of the seven mental processes using an independent Chinese Human Connectome Project (CHCP) of 228 subjects. By retraining the model parameters for thirty times using the same strategy on the CHCP dataset, the functional neuroanatomy fingerprint also shows satisfying classification performance ability area under the curve (AUC) of 0.99 among the seven mental processes (Fig. 4A). In a more stringent situation where we directly apply the model trained on HCP dataset without any parameter updates to classify CHCP dataset, we still obtain satisfying mean classification accuracy of 86% (Fig. 4C), and vice versa. Moreover, the identified functional neuroanatomy fingerprint using CHCP dataset is robust in terms of low standard deviation of functional activity weight value in each region visualized in Fig. 4B, and similar to those using HCP dataset (Supplemental Figs. 2-4) by calculating the Spearman correlation coefficients of the two fingerprints using HCP and CHCP respectively in each of the seven mental processes (Fig. 5). By grouping the whole-brain neuroanatomical regions into the six categories including five cortical lobes and one subcortical area, the occipital lobe also consistently exhibits significantly larger ‘importance’ than the other five categories (six-way ANOVA, p<0.001, FDR corrected, post-hoc t-test) in most mental processes except for Language and Motor under different thresholds of top *k%* important regions (Fig. 4D and Supplemental Fig. 5) across the two independent datasets. Moreover, similar as in Fig. 3D, sulci consistently exhibit significantly larger ‘importance’ than gyri and subcortical area (three-way ANOVA, p<0.001, FDR corrected, post-hoc t-test) under different thresholds of top *k%* important regions and across all seven mental processes in CHCP dataset as illustrated in Fig. 4E and Supplemental Figs. 6-7.

**Fig. 4.**
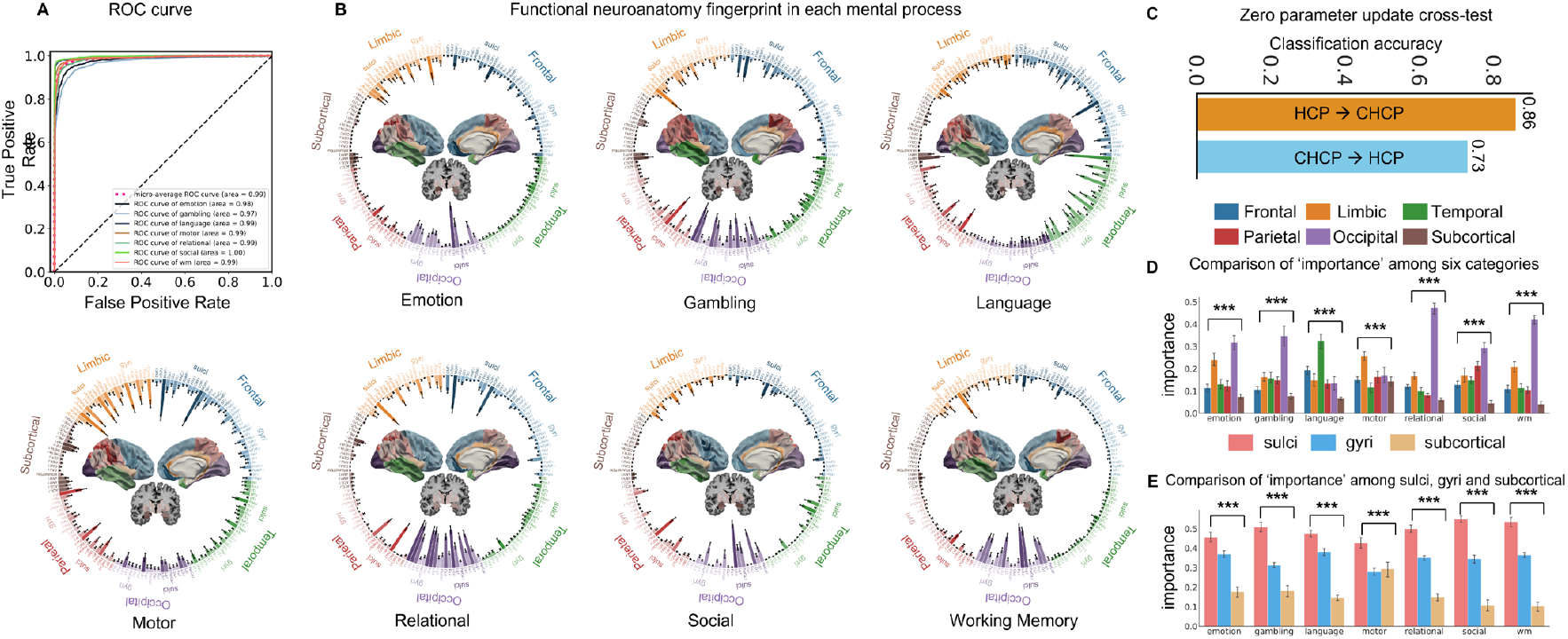
Robustness of functional neuroanatomy fingerprint in the independent CHCP dataset. **(A)** ROC curves for each of the seven mental processes. **(B)** Representation of the functional neuroanatomy fingerprint in each mental process as a set of functional activity weight values of all neuroanatomical regions defined by the Desikan-Killiany (DK) brain atlas. Note that each value is represented as mean±std based on the thirty trained models. **(C)** Zero parameter update of classification between the two datasets. We directly apply the model trained on HCP dataset without any parameter updates to classify CHCP dataset,and vice versa. **(D)** Comparison of ‘importance’ among the six categories (five cortical lobes and one subcortical area) in each mental process using top 30% of important brain regions. **(E)** Comparison of ‘importance’ value among gyri, sulci, and subcortical area in each of the mental process using top 30% of important brain regions. The data is represented as the mean value with 95% confidence intervals, *** indicates p<0.001 in multi-way ANOVA. FDR correction is used for multiple comparisons.

**Fig. 5.**
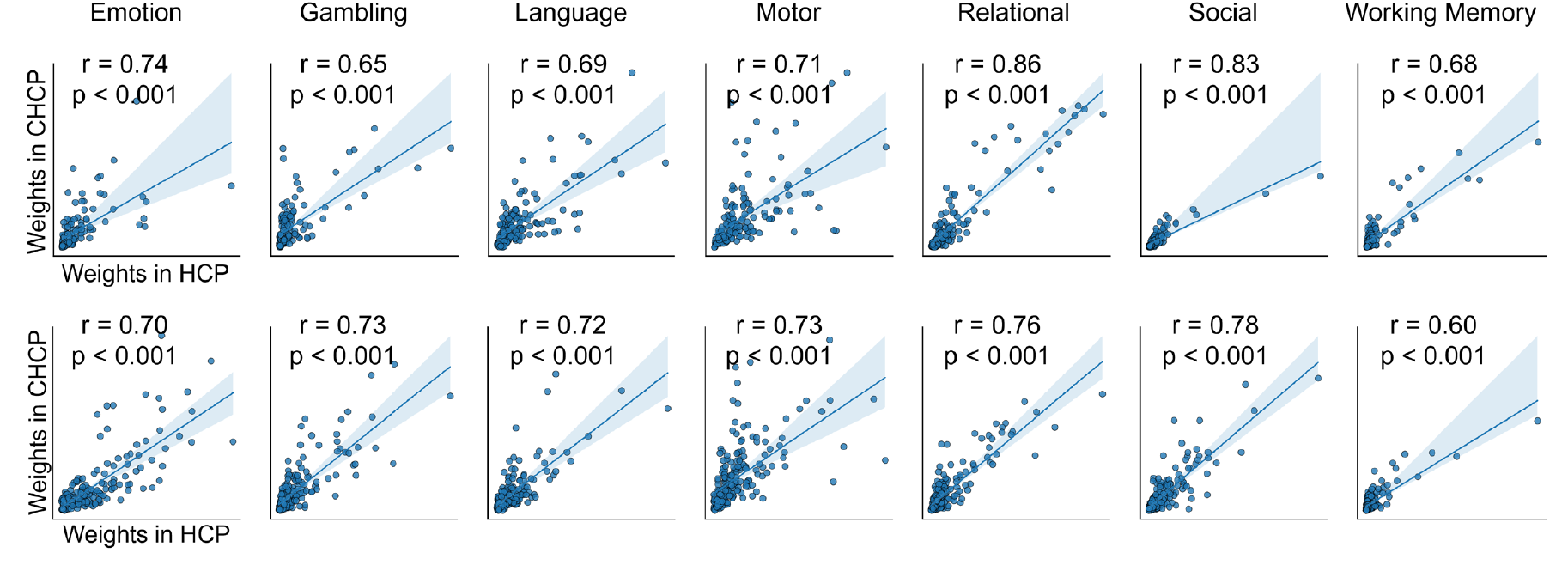
Similarity of functional neuroanatomy fingerprint between HCP and CHCP datasets. In each sub-figure, the horizontal and vertical axis represents the weights within the fingerprints using HCP and CHCP, respectively. The similarity is calculated by the Spearman correlation coefficient of the two fingerprints between HCP and CHCP. The first and second rows represent the similarity of fingerprints based on the DK and Schaefer brain atlases, respectively.

We further assess the robustness of identified functional neuroanatomy fingerprint using another Schaefer brain atlas. The visualizations of the identified functional neuroanatomy fingerprint in the Schaefer brain atlas are in Supplemental Figs. 7-12, and achieve satisfying mean classification accuracy of 82% when performing zero parameter update of classification between the two independent datasets (Supplemental Fig. 13). Since the Schaefer atlas has finer-grained neuroanatomical regions (100 gyri/sulci regions plus 19 subcortical regions) compared to the DK atlas, we group all regions into six categories including five cortical lobes (frontal, parietal, temporal, occipital, and limbic) and one subcortical area for a fair comparison of fingerprints between the two atlases. The distribution pattern of functional activity weights in the identified functional neuroanatomy fingerprint using DK atlas is similar to that using Schaefer brain atlas (Supplemental Figs. 3-4). Fig. 6 shows that similar to that using DK atlas (Fig. 1), the occipital lobe exhibits significantly larger ‘importance’ than the other five categories (six-way ANOVA, p<0.001, FDR corrected, post-hoc t-test) in most mental processes except for Language and Motor across different thresholds of top *k%* important regions.

**Fig. 6.**
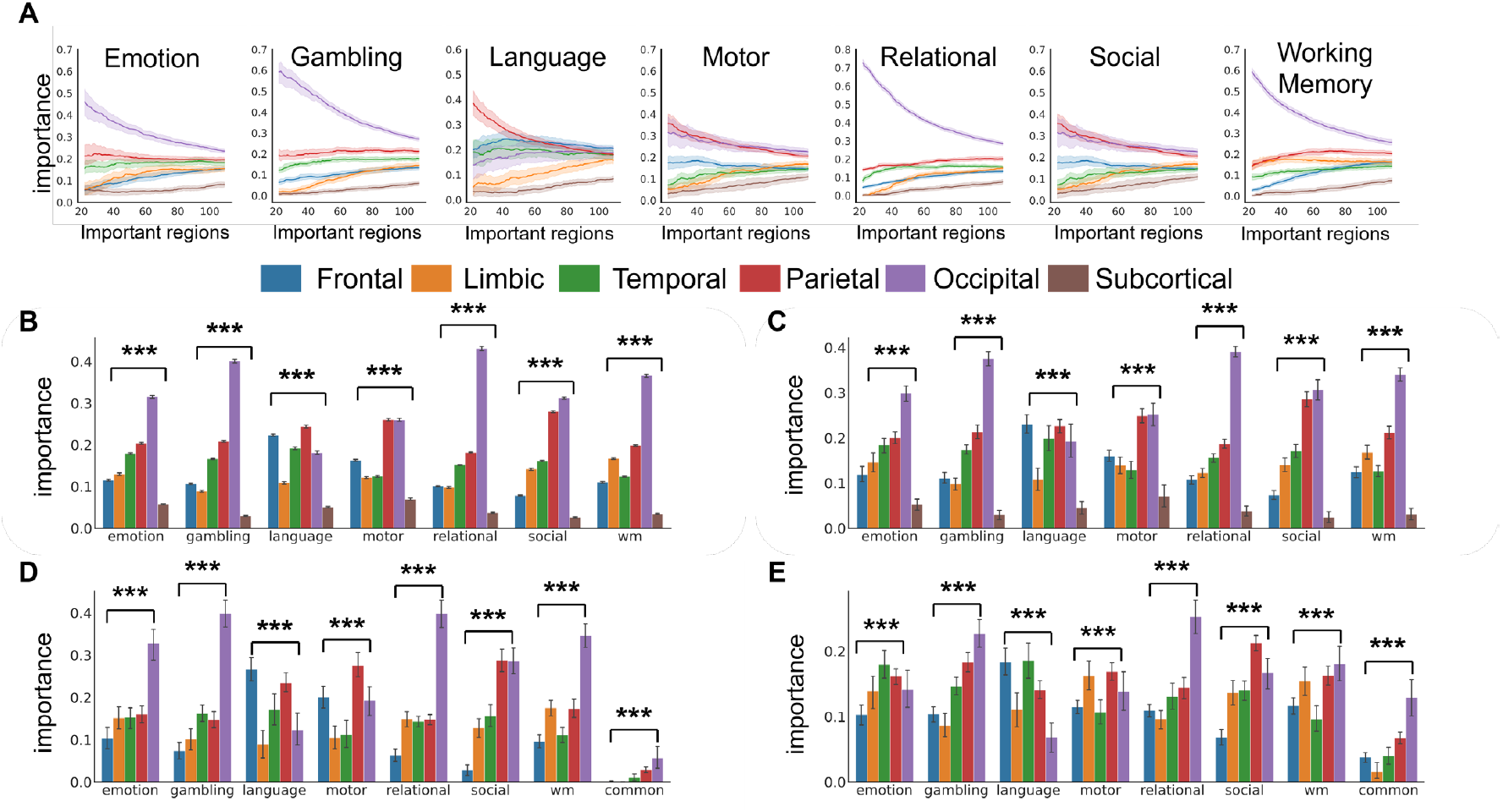
Comparisons of ‘importance’ among the six categories of fingerprints in HCP dataset using the Schaefer atlas. **(A)** Comparison of ‘importance’ among the six categories in each mental process under different thresholds of important brain regions from top 10% (16 brain regions) to top 50% (80 brain regions) with a 0.5% interval. The data is represented as the mean value with 95% confidence intervals. **(B)** Statistical comparison of ‘importance’ among the six categories in each mental process in (A). **(C)** Comparison of ‘importance’ among the six categories in each mental process using top 30% of important brain regions. **(D)** Comparison of ‘importance’ of gyri among the five cortical lobes in the unique and common regions, respectively using top 30% of important brain regions. **(E)** Comparison of ‘importance’ of sulci among the five cortical lobes in the unique and common regions, respectively using top 30% of important brain regions. The data is represented as the mean value with 95% confidence intervals, *** indicates p<0.001 in multi-way ANOVA. FDR correction is used for multiple comparisons.

## Discussion and Conclusion

In this study, we proposed a multi-task machine learning framework based on task fMRI data to address a fundamental question: whether a specific mental process possesses a ‘functional neuroanatomy fingerprint’ which refers to a unique and reliable pattern of functional neuroanatomy to underlie how the mental process receives, encodes, stores, retrieves, and manipulates information in the brain. Using the task fMRI data of seven representative and different mental processes including Emotion, Gambling, Language, Motor, Relational, Social, and Working Memory across two independent cohorts of 1235 subjects from the US and China as a test bed, experimental results demonstrated that our proposed framework can effectively disentangle the functional neuroanatomy fingerprint of the seven mental processes, each of which was characterized by a unique set of functional activity weights of whole-brain regions. The identified functional neuroanatomy fingerprint of each mental process was consistent across two independent datasets and two different brain atlases, and achieved satisfying discriminative ability of 93% classification accuracy and AUC of 0.99 from those of the other mental processes. This study provided one of the earliest neuroimaging-based evidences of the functional neuroanatomy substrates for mental processing.

By investigating the commonality and uniqueness of the fingerprints across seven mental processes, we found that occipital lobe, parietal lobe, and amygdala exhibited common high degree of involvement across all seven fingerprints. Amygdala emerged as an important subcortical region across different processes due to its various functions in emotion processing, memory, social cognition, etc. (*61-66*). The uniqueness of each fingerprint was mainly reflected by the strong association with the corresponding task paradigm of the mental process, i.e., the temporal lobe responsible for language processing showed statistically larger functional activity weights within the fingerprint of Language compared to those of the other mental processes.

Previous studies have proposed various hypotheses of brain cortical folding into gyri and sulci such as brain skull restriction (*38,39*), differential laminar growth (*40*), genetic regulation (*41*), and axonal tension or pulling (*42,43*). While the exact neuromechanism of cortical folding remains largely unknown, researchers have extensively reported the significant differences between gyri and sulci in terms of anatomy (*44,45*), morphology (*46*), functional property (*17,47,48*), structural property (*49*), as well as differential association with certain mental processes (*47*). In this study, we divided each cortical region into fine-scale gyri and sulci, and systematically investigated the differential degree of involvement between gyri and sulci in the identified functional neuroanatomy fingerprint of each mental process. We found that the gyri and sulci of lateral occipital (LO) region were both involved in all of the seven fingerprints. LO is responsible for visual stimuli formation, progressing from local to global perception (*50,51*). It combines the functions of primary vision formation (concrete) and higher vision formation (abstract) (*52*). In most of other regions including pericalcarine, fusiform, lingual regions, superior parietal region, and orbital frontal region, sulci were more involved than gyri in the fingerprint. Pericalcarine, fusiform, and lingual regions are specialized in higher-order visual processing such as face recognition (*53-55*), superior parietal region is primarily responsible for motor-sensory integration (*56*), orbital frontal region is primarily associated with emotion processing (*57*). In limbic lobe, gyri were more involved than sulci. These findings highlighted differential degree of involvement between gyri and sulci in mental processing. Especially, sulci were more involved in higher-order and specialized functions, indicating its crucial role in discriminating different mental processes. The different roles between gyri and sulci in discriminating the functional neuroanatomy fingerprint of each mental process are further supported by previous studies demonstrating that sulci mainly serve as local functional processing units, while gyri are mainly responsible for global functional information exchange (*47*). Specifically, sulci exhibited weaker long-range functional connectivity strength (*49*) and stronger magnitude of power spectral density of fMRI signal (*58*) compared to gyri. Moreover, gyri play a more important role in discriminating task state and resting state (*17*), while sulci are more crucial in discriminating different task states (i.e., mental processing) (*44,59,60*).

In conclusion, this study combined both advance machine learning techniques and large cohorts of neuroimaging data to provide a solid functional neuroanatomy foundation to enhance our understanding of the neural mechanism of mental processing. One limitation of this study was that given the available data, we merely used seven task-based fMRI datasets of young adult subjects in two independent datasets as a test bed for identification and characterization of the functional neuroanatomy fingerprint of a specific mental process. More task-based fMRI data representing other mental processes in a wider age range group are needed to further confirm that all mental processes possess a unique and reliable functional neuroanatomy fingerprint. In addition, the regularity and variability of identified functional neuroanatomy fingerprint of a specific mental process across different age groups, different cultures, or even different psychiatric disorders (*80-81*) are also promising future work.

## Materials and Methods

### Study Overview

The overview of this study (Fig. 7) mainly includes three parts: data preprocessing (Fig. 7A), identification of functional neuroanatomy fingerprint (Fig. 7B), and interpretation and validation of the functional neuroanatomy fingerprint (Fig. 7C). We introduce the three parts in details in the following sections.

**Fig. 7.**
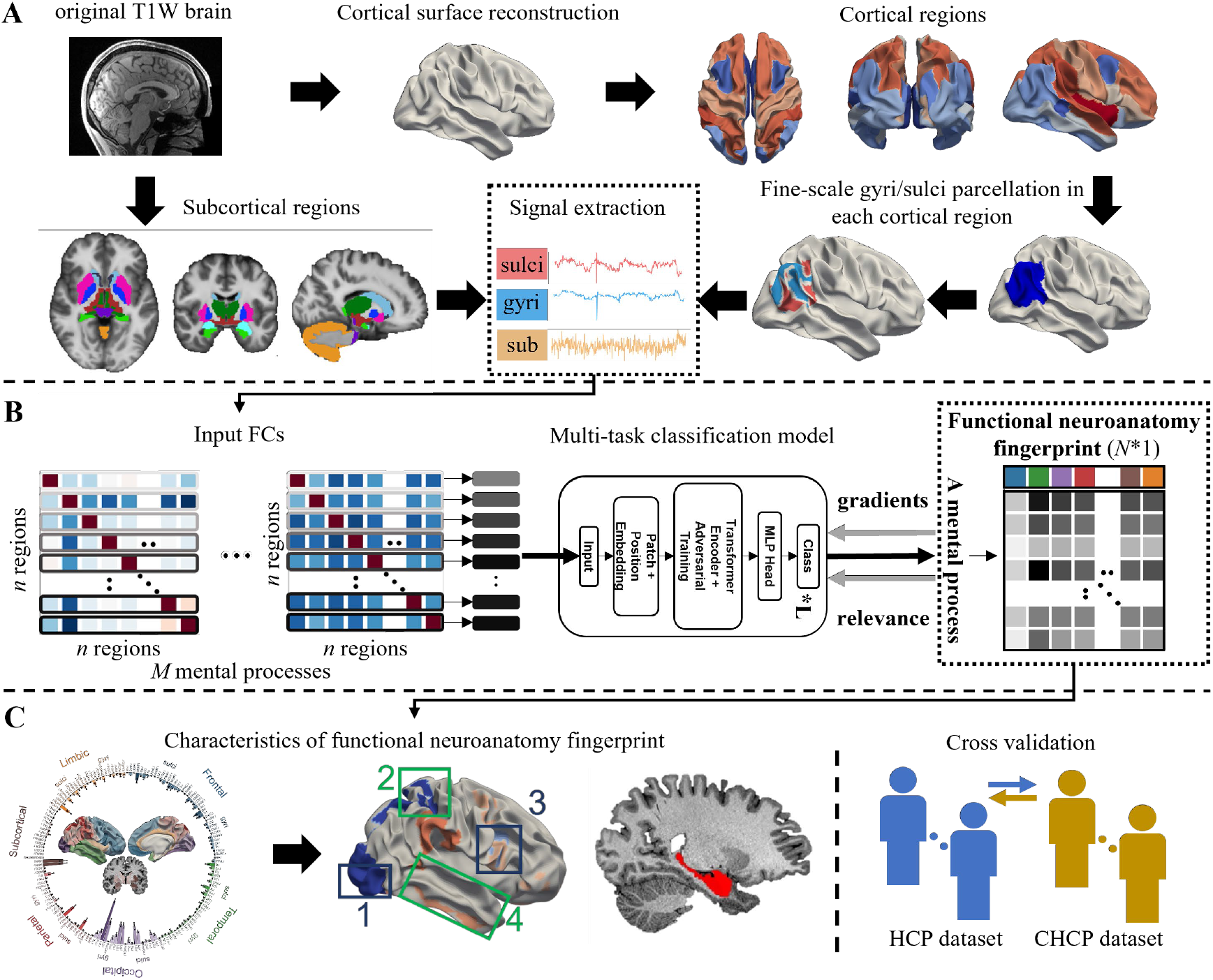
Overview of this study. **(A)** Data preprocessing. The data preprocessing mainly includes cortical surface reconstruction based on raw MRI data, parcellation of cortical and subcortical brain regions based on brain atlases, fine-scale gyri/sulci parcellation of each cortical region, and fMRI signal extraction of each gyral, sulcal, and subcortical brain region. **(B)** Identification of functional neuroanatomy fingerprint. The whole-brain functional connectivity (FC) matrix is calculated in each mental process of each subject based on the extracted fMRI signals, and is used as the model input. The proposed multi-task deep learning classification model outputs the multi-class classification label as well as the associated fingerprint contributing to the classification. **(C)** Characterization of the functional neuroanatomy fingerprint. The characteristics of the identified fingerprint such as the distribution and importance of involved neuroanatomy regions is assessed. The robustness of identified fingerprint is also validated using different datasets and brain atlases.

### Data Description and Preprocessing

Two independent datasets from the US and China were used in this study: Human Connectome Project (HCP) (*67-70*) and Chinese Human Connectome Project (CHCP) (*71,72*). Both of the two datasets contain task-based functional MRI (t-fMRI) data of the same seven task designs representing seven mental processes, facilitating the evaluation of robustness of functional neuroanatomy fingerprint across the two independent datasets. The seven mental processes cover a wide range of neural systems including 1) emotion processing; 2) category specific representations; 3) language processing; 4) visual, motion, somatosensory, and motor systems; 5) relational processing; 6) social cognition; and 7) working memory/cognitive control systems, and are abbreviated to Emotion, Gambling, Language, Motor, Relational, Social, and Working Memory, respectively. The detailed task design descriptions of each mental process are referred to (*71-73*).

A total of 1,007 subjects (22-35 years old with mean age of 28.72 ± 3.69 years) having t-fMRI data of all seven mental processes were adopted in the HCP dataset. All subjects were scanned using a customized Siemens 3T “Connectome Skyra” with a standard 32-channel Siemens receive head coil at Washington University in St. Louis, the US. The major scanning parameters of structural MRI T1w data are: repetition time = 24,000 ms, flip angle = 8^°^, field of view = 224 × 224 mm, voxel size = 0.7 mm isotropic, bandwidth = 210 Hz/Px. The t-fMRI data were acquired with the following major parameters: repetition time = 720 ms, echo time = 33.1 ms, flip angle = 52^°^, field of view = 208 × 180 mm, matrix = 104 × 94, slice thickness = 2.0 mm, voxel size = 2.0 mm isotropic, multiband factor = 8, bandwidth = 2290 Hz/Px. A total of 228 subjects (18-31 years old with mean age of 22.28 ± 2.84 years) with the same all seven t-fMRI data were adopted in the CHCP dataset. All subjects were scanned using a Siemens 3T Prisma scanner in Beijing, China. The major acquisition parameters of structural MRI T1w are: repetition time = 24,000 ms, echo time = 2.22 ms, flip angle = 8^°^, field of view = 256 × 240 mm, voxel size = 0.8 mm isotropic, bandwidth = 220 Hz/Px. The major parameters of t-fMRI data are: repetition time = 710 ms, echo time = 33.0 ms, flip angle = 52^°^, field of view = 212 × 212 mm, matrix = 104 × 94, slice thickness = 2.0 mm, voxel size = 2.0 mm isotropic, multiband factor = 8, bandwidth = 2358 Hz/Px.

The minimal preprocessing pipeline (*74*, https://github.com/Washington-University/HCPpipelines) were consistently applied to the raw MRI data of the two datasets. Each individual brain was parcellated into 19 subcortical regions based on the FreeSurfer atlas as well as 68/100 cortical regions based on the Desikan-Killiany (DK)/Schaefer cortical atlas to evaluate the robustness of the fingerprint across different brain atlases. Considering the importance of brain cortical folding in mental processing (*38-43*) as well as the functional property differentiation between cortical gyri and sulci (*44-49*), we further divided each cortical region into fine-scale gyral and sulcal areas based on the sulc value (*75-76*), resulting 19+68*2=155 or 19+100*2=219 brain regions. We finally extracted the mean preprocessed t-fMRI signal of each parcellated cortical and subcortical brain region, and calculated the Pearson Correlation Coefficient of two t-fMRI signals between any pair of brain regions to obtain the symmetric functional connectivity matrix.

### Multi-Class Classification Model of Mental Processes

The calculated functional connectivity (FC) matrices of each of the seven mental processes in each subject were used as the input of multi-class classification model to differentiate different mental processes. The proposed multi-class classification model is based on the Vision Transformer (ViT) (*35*) mainly including embedding (patch and position), transformer blocks, and a fully connected layer (Fig. 7B). The model outputs the multi-class classification label as well as the associated fingerprint contributing to the classification.

Specifically, the input symmetric FC matrix *x* ∈ℝ^*N∗N*^(*N* is the number of brain regions) was converted into *N* nonoverlapped patches *x*_*p*_ ∈ ℝ^1*∗N*^. Unlike the traditional patch size such as 16*16 or 14*14 in natural image processing, each row of the FC representing the functional connectivity characteristics of one brain region to the other ones was defined as a meaningful patch. We then put each patch into a trainable linear projection layer to obtain *x*_*pa*_ ∈ ℝ^1*∗D*^(*D* denotes the target dimension and was set to 784). The positional embedding *x*_*po*_ ∈ ℝ^*N∗D*^ was introduced to retain positional information. We also added an extra class token *Z*_*cls*_ ∈ ℝ^1*∗D*^ before the patch embedding. The concatenation of both patch and positional embeddings *x*_*pec*_ ∈ ℝ ^(*N*+1)*∗D*^ was used as input to the transformer block. The transformer block consisted of one Multi-head Self-Attention (MSA) block and one Multi-Layer Perception (MLP) block. Each block was preceded by a layer normalization (LayerNorm). In the MSA layer, we used *K* heads to divide the input *x* ∈ *ℝ* ^(*N*+1)*∗D*^ into 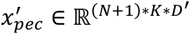, where 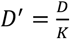. After performing the attention calculation (*35*), the outputs of all *K* heads were then concatenated and fed into the MLP layer to output class token *Z*_*cls*_. In this study, we used 12 transformer blocks and an MLP for classification, and performed five-fold cross-validation to obtain the averaged classification result. The detailed model parameter settings are provided in Supplemental Tables 1-2. Moreover, we incorporated adversarial training into the multi-class classification model to increase both robustness (*77, 78*) and interpretability (*79*) of the model. The Fast Gradient Sign Method (FGSM) (*78*) was adopted to generate adversarial samples.

### Identification and Characterization of Functional Neuroanatomy Fingerprint

Based on the trained multi-class classification model of t-fMRI-derived FCs of seven mental processes, we identified the functional neuroanatomy fingerprint of each mental process. Specifically, we adopted an attribution propagation method named Layer-wise Relevance Propagation (LRP) with a class-specific approach to obtain the attention map of a specific mental process (*36*). Within the first row of the attention matrix, the magnitude of each element reflected the degree of similarity or contribution between each patch (i.e., brain region) and the class token (i.e., a mental process). Therefore, the first row of attention matrix consisting of a set of functional activity weight values of all neuroanatomical regions was defined as the functional neuroanatomy fingerprint of the mental process (Fig. 7B).

We characterized the identified functional neuroanatomy fingerprint of the mental process from several aspects (Fig. 7C). We first defined top *k*% regions with largest functional activity weight values as the ‘important regions’ contributing to the fingerprint, and then assessed and compared the ‘importance’ value as normalized ratio of important regions among the six brain region categories (five lobes and one subcortical area) or among cortical gyri, cortical sulci, and subcortical regions within a specific mental process. The commonality and uniqueness of such ‘importance’ distribution among the six categories or among gyri, sulci, and subcortical were also compared among the seven mental processes. Finally, the robustness of identified functional neuroanatomy fingerprint as well as its characteristics were validated using two independent datasets (HCP and CHCP) and two different brain cortical atlases (DK and Schaefer).

## Acknowledgments

Data were provided in part by the Chinese Human Connectome Project (CHCP, PI: Jia-Hong Gao) funded by the Beijing Municipal Science & Technology Commission, Chinese Institute for Brain Research (Beijing), National Natural Science Foundation of China, and the Ministry of Science and Technology of China. Data were provided in part by the Human Connectome Project, WU-Minn Consortium (Principal Investigators: David Van Essen and Kamil Ugurbil; 1U54MH091657) funded by the 16 NIH Institutes and Centers that support the NIH Blueprint for Neuroscience Research; and by the McDonnell Center for Systems Neuroscience at Washington University.

## Author Contributions

Conceptualization: Zifan Wang, Xi Jiang; Methodology: Zifan Wang, Yuzhong Chen, Wei Mao, Zhenxiang Xiao; Visualization: Zifan Wang, Xi Jiang; Funding acquisition: Xi Jiang; Project administration: Xi Jiang, Alan C Evans; Data curation: Zifan Wang, Guannan Cao, Zhenxiang Xiao, Tuo Zhang; Supervision: Xi Jiang, Alan C Evans, Paule-J Toussaint; Writing – original draft: Zifan Wang; Writing – review & editing: Xi Jiang, Jiadong Yan, Yi Pan, Paule-J Toussaint, Alan C Evans.

## Data and Code Availability

The HCP raw dataset is available in https://www.humanconnectome.org/. The CHCP raw dataset is available in https://www.Chinese-HCP.cn. Part of the preprocessed data and results are available upon request. The code of this study is available on the GitHub (https://github.com/XiJiangLabUESTC/Fingerprint).

## References

1. W. R. Uttal, The new phrenology: The limits of localizing cognitive processes in the brain. (The MIT press, 2001).

2. R. J. Joynt, Paul Pierre Broca: His contribution to the knowledge of aphasia. Cortex 1, 206–213 (1964).

3. W. Penfield, E. Boldrey, Somatic motor and sensory representation in the cerebral cortex of man as studied by electrical stimulation. Brain 60, 389–443 (1937).

4. K. Brodmann, Vergleichende Lokalisationslehre der Grosshirnrinde in ihren Prinzipien dargestellt auf Grund des Zellenbaues. (Barth, 1909).

5. L. R. Squire, J. T. Wixted, The cognitive neuroscience of human memory since HM. Annual review of neuroscience 34, 259–288 (2011).

6. H. Klüver, P. C. Bucy, “ Psychic blindness” and other symptoms following bilateral temporal lobectomy in Rhesus monkeys. American Journal of Physiology, (1937).

7. R. S. Desikan et al., An automated labeling system for subdividing the human cerebral cortex on MRI scans into gyral based regions of interest. Neuroimage 31, 968–980 (2006).

8. A. Schaefer et al., Local-global parcellation of the human cerebral cortex from intrinsic functional connectivity MRI. Cerebral cortex 28, 3095–3114 (2018).

9. G. Tononi, O. Sporns, G. M. Edelman, A measure for brain complexity: relating functional segregation and integration in the nervous system. Proceedings of the National Academy of Sciences 91, 5033–5037 (1994).

10. K. Friston, Beyond phrenology: what can neuroimaging tell us about distributed circuitry? Annual review of neuroscience 25, 221–250 (2002).

11. Y. Du et al., NeuroMark: An automated and adaptive ICA based pipeline to identify reproducible fMRI markers of brain disorders. NeuroImage: Clinical 28, 102375 (2020).

12. K. J. Friston, Functional and effective connectivity: a review. Brain connectivity 1, 13–36 (2011).

13. K. J. Friston, Modalities, Modes, and Models in Functional Neuroimaging. Science 326, 399–403 (2009).

14. J. Duncan, The multiple-demand (MD) system of the primate brain: mental programs for intelligent behaviour. Trends in cognitive sciences 14, 172–179 (2010).

15. M. W. Cole, W. Schneider, The cognitive control network: Integrated cortical regions with dissociable functions. Neuroimage 37, 343–360 (2007).

16. B. T. T. Yeo et al., The organization of the human cerebral cortex estimated by intrinsic functional connectivity. Journal of neurophysiology, (2011).

17. M. Jiang et al., Anatomy-guided spatio-temporal graph convolutional networks (AG-STGCNs) for modeling functional connectivity between gyri and sulci across multiple task domains. IEEE Transactions on Neural Networks and Learning Systems, (2022).

18. M. Schurz, L. Maliske, P. Kanske, Cross-network interactions in social cognition: A review of findings on task related brain activation and connectivity. cortex 130, 142–157 (2020).

19. R. Jiang et al., Task-induced brain connectivity promotes the detection of individual differences in brain-behavior relationships. NeuroImage 207, 116370 (2020).

20. A. Segal et al., Regional, circuit and network heterogeneity of brain abnormalities in psychiatric disorders. Nature Neuroscience, 1-17 (2023).

21. S. Han, Y. Ma, Cultural differences in human brain activity: A quantitative meta-analysis. NeuroImage 99, 293–300 (2014).

22. Y. Zhu, L. Zhang, J. Fan, S. Han, Neural basis of cultural influence on self-representation. NeuroImage 34, 1310–1316 (2007).

23. S. Han, G. Northoff, Culture-sensitive neural substrates of human cognition: a transcultural neuroimaging approach. Nature Reviews Neuroscience 9, 646–654 (2008).

24. T. Harada et al., Cultural influences on neural systems of intergroup emotion perception: An fMRI study. Neuropsychologia 137, 107254 (2020).

25. S. J. Morrison, S. M. Demorest, E. H. Aylward, S. C. Cramer, K. R. Maravilla, FMRI investigation of cross-cultural music comprehension. NeuroImage 20, 378–384 (2003).

26. B. Biswal, F. Zerrin Yetkin, V. M. Haughton, J. S. Hyde, Functional connectivity in the motor cortex of resting human brain using echo-planar MRI. Magnetic resonance in medicine 34, 537–541 (1995).

27. M. M. Monti, Statistical analysis of fMRI time-series: a critical review of the GLM approach. Frontiers in human neuroscience 5, 28 (2011).

28. M. J. McKeown et al., Spatially independent activity patterns in functional MRI data during the Stroop color-naming task. Proceedings of the National Academy of Sciences 95, 803–810 (1998).

29. R. Viviani, G. Grön, M. Spitzer, Functional principal component analysis of fMRI data. Human brain mapping 24, 109–129 (2005).

30. J. Lv et al., Sparse representation of whole-brain fMRI signals for identification of functional networks. Medical image analysis 20, 112–134 (2015).

31. X. Wang et al., Decoding and mapping task states of the human brain via deep learning. Human brain mapping 41, 1505–1519 (2020).

32. B. P. Rogers, V. L. Morgan, A. T. Newton, J. C. Gore, Assessing functional connectivity in the human brain by fMRI. Magnetic resonance imaging 25, 1347–1357 (2007).

33. E. S. Finn et al., Functional connectome fingerprinting: identifying individuals using patterns of brain connectivity. Nature Neuroscience 18, 1664–1671 (2015).

34. J. Gonzalez-Castillo et al., Whole-brain, time-locked activation with simple tasks revealed using massive averaging and model-free analysis. Proceedings of the National Academy of Sciences 109, 5487–5492 (2012).

35. A. Dosovitskiy et al., An image is worth 16x16 words: Transformers for image recognition at scale. arXiv preprint 2010.11929, (2020).

36. H. Chefer, S. Gur, L. Wolf. pp. 782–791.

37. D. M. Barch et al., Function in the human connectome: task-fMRI and individual differences in behavior. Neuroimage 80, 169–189 (2013).

38. W. Welker, Why does cerebral cortex fissure and fold? A review of determinants of gyri and sulci. Cerebral Cortex: comparative structure and evolution of Cerebral Cortex, Part II, 3-136 (1990).

39. E. A. Miska et al., Microarray analysis of microRNA expression in the developing mammalian brain. Genome biology 5, 1–13 (2004).

40. D. P. Richman, R. M. Stewart, J. Hutchinson, V. S. Caviness Jr, Mechanical Model of Brain Convolutional Development: Pathologic and experimental data suggest a model based on differential growth within the cerebral cortex. Science 189, 18–21 (1975).

41. B. Mota, S. Herculano-Houzel, How the cortex gets its folds: an inside-out, connectivity-driven model for the scaling of mammalian cortical folding. Frontiers in neuroanatomy 6, 3 (2012).

42. D. C. v. Essen, A tension-based theory of morphogenesis and compact wiring in the central nervous system. Nature 385, 313–318 (1997).

43. J. Nie et al., Axonal fiber terminations concentrate on gyri. Cerebral cortex 22, 2831–2839 (2012).

44. I. H. M. Smart, C. Dehay, P. Giroud, M. Berland, H. Kennedy, Unique morphological features of the proliferative zones and postmitotic compartments of the neural epithelium giving rise to striate and extrastriate cortex in the monkey. Cerebral cortex 12, 37–53 (2002).

45. B. Fischl, A. M. Dale, Measuring the thickness of the human cerebral cortex from magnetic resonance images. Proceedings of the National Academy of Sciences 97, 11050–11055 (2000).

46. V. A. Magnotta et al., Quantitative in vivo measurement of gyrification in the human brain: changes associated with aging. Cerebral Cortex 9, 151–160 (1999).

47. X. Jiang, T. Zhang, S. Zhang, K. M. Kendrick, T. Liu, Fundamental functional differences between gyri and sulci: implications for brain function, cognition, and behavior. Psychoradiology 1, 23–41 (2021).

48. H. Liu et al., The cerebral cortex is bisectionally segregated into two fundamentally different functional units of gyri and sulci. Cerebral Cortex 29, 4238–4252 (2019).

49. F. Deng et al., A functional model of cortical gyri and sulci. Brain structure and function 219, 1473–1491 (2014).

50. R. Malach et al., Object-related activity revealed by functional magnetic resonance imaging in human occipital cortex. Proceedings of the National Academy of Sciences 92, 8135–8139 (1995).

51. K. Grill-Spector et al., Differential processing of objects under various viewing conditions in the human lateral occipital complex. Neuron 24, 187–203 (1999).

52. Z. Kourtzi, N. Kanwisher, Representation of perceived object shape by the human lateral occipital complex. Science 293, 1506–1509 (2001).

53. N. Kanwisher, J. McDermott, M. M. Chun, The fusiform face area: a module in human extrastriate cortex specialized for face perception. Journal of neuroscience 17, 4302–4311 (1997).

54. N. Kanwisher, G. Yovel, The fusiform face area: a cortical region specialized for the perception of faces. Philosophical Transactions of the Royal Society B: Biological Sciences 361, 2109–2128 (2006).

55. A. Mechelli, G. W. Humphreys, K. Mayall, A. Olson, C. J. Price, Differential effects of word length and visual contrast in the fusiform and lingual gyri during. Proceedings of the Royal Society of London. Series B: Biological Sciences 267, 1909–1913 (2000).

56. D. M. Wolpert, S. J. Goodbody, M. Husain, Maintaining internal representations: the role of the human superior parietal lobe. Nature neuroscience 1, 529–533 (1998).

57. C. A. Hynes, A. A. Baird, S. T. Grafton, Differential role of the orbital frontal lobe in emotional versus cognitive perspective-taking. Neuropsychologia 44, 374–383 (2006).

58. S. Zhang et al., Deep Learning Models Unveiled Functional Difference Between Cortical Gyri and Sulci. IEEE Trans Biomed Eng 66, 1297–1308 (2019).

59. A. Fornito et al., Individual differences in anterior cingulate/paracingulate morphology are related to executive functions in healthy males. Cerebral cortex 14, 424–431 (2004).

60. A. Cachia et al., How interindividual differences in brain anatomy shape reading accuracy. Brain Structure and Function 223, 701–712 (2018).

61. L. W. Swanson, G. D. Petrovich, What is the amygdala? Trends in neurosciences 21, 323–331 (1998).

62. P. H. Janak, K. M. Tye, From circuits to behaviour in the amygdala. Nature 517, 284–292 (2015).

63. J. E. Ledoux, Emotion and the amygdala. (1992).

64. J. LeDoux, The amygdala. Current biology 17, R868–R874 (2007).

65. G. Z. Tau, B. S. Peterson, Normal development of brain circuits. Neuropsychopharmacology 35, 147–168 (2010).

66. M. A. Sommer, R. H. Wurtz, Brain circuits for the internal monitoring of movements. Annu. Rev. Neurosci. 31, 317–338 (2008).

67. D. C. Van Essen et al., The WU-Minn human connectome project: an overview. Neuroimage 80, 62–79 (2013).

68. M. F. Glasser et al., The minimal preprocessing pipelines for the Human Connectome Project. Neuroimage 80, 105–124 (2013).

69. J. S. Elam et al., The human connectome project: a retrospective. NeuroImage 244, 118543 (2021).

70. D. C. Van Essen et al., The Human Connectome Project: a data acquisition perspective. Neuroimage 62, 2222–2231 (2012).

71. J. Ge et al., Increasing diversity in connectomics with the Chinese Human Connectome Project. Nature Neuroscience 26, 163–172 (2023).

72. N. Vogt, The Chinese Human Connectome Project. Nature Methods 20, 177–177 (2023).

73. D. M. Barch et al., Function in the human connectome: task-fMRI and individual differences in behavior. Neuroimage 80, 169–189 (2013).

74. M. F. Glasser et al., The minimal preprocessing pipelines for the Human Connectome Project. 80, 105–124 (2013).

75. B. Fischl, M. I. Sereno, A. M. Dale, Cortical surface-based analysis: II: inflation, flattening, and a surface-based coordinate system. Neuroimage 9, 195–207 (1999).

76. S. Yang et al., Temporal variability of cortical gyral-sulcal resting state functional activity correlates with fluid intelligence. Frontiers in neural circuits 13, 36 (2019).

77. T. Miyato, A. M. Dai, I. Goodfellow, Adversarial training methods for semi-supervised text classification. arXiv preprint 1605.07725, (2016).

78. I. J. Goodfellow, J. Shlens, C. Szegedy, Explaining and harnessing adversarial examples. arXiv preprint 1412.6572, (2014).

79. T. Zhang, Z. Zhu. (PMLR), pp. 7502–7511.

80. X. Jiang, X.J. Shou, Z. Zhao, Y. Chen, F.C. Meng, J. Le, … & R. Zhang A brain structural connectivity biomarker for autism spectrum disorder diagnosis in early childhood, Psychoradiology, 3:kkad005. (2023)

81. J. Sui, D. Zhi, & V.D. Calhoun. Data-driven multimodal fusion: approaches and applications in psychiatric research. Psychoradiology, 3, kkad026. 10.1093/psyrad/kkad026. (2023)

