## Supplementary Info for "Multi-task machine learning reveals the functional neuroanatomy fingerprint of mental processing" (from another version of this preprint: version 2, 2025-07-03)

**One-Sentence Summary:** There exists a functional neuroanatomy fingerprint to underlie the mental process.

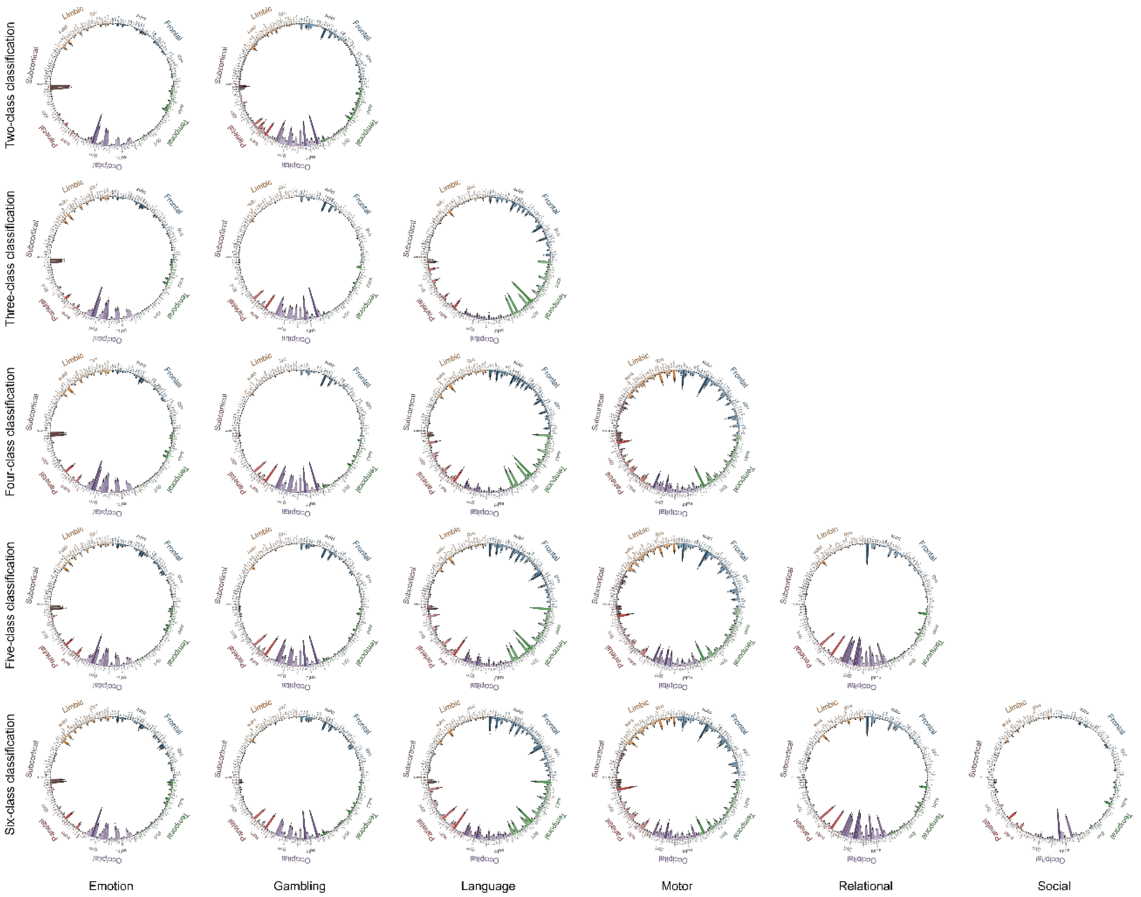


**Supplemental Fig. 1 Stability of functional neuroanatomy fingerprint under different types of classification models.** The fingerprint of Emotion is used as a benchmark for stability assessment from two-class classification to six-class classification with other mental processes.

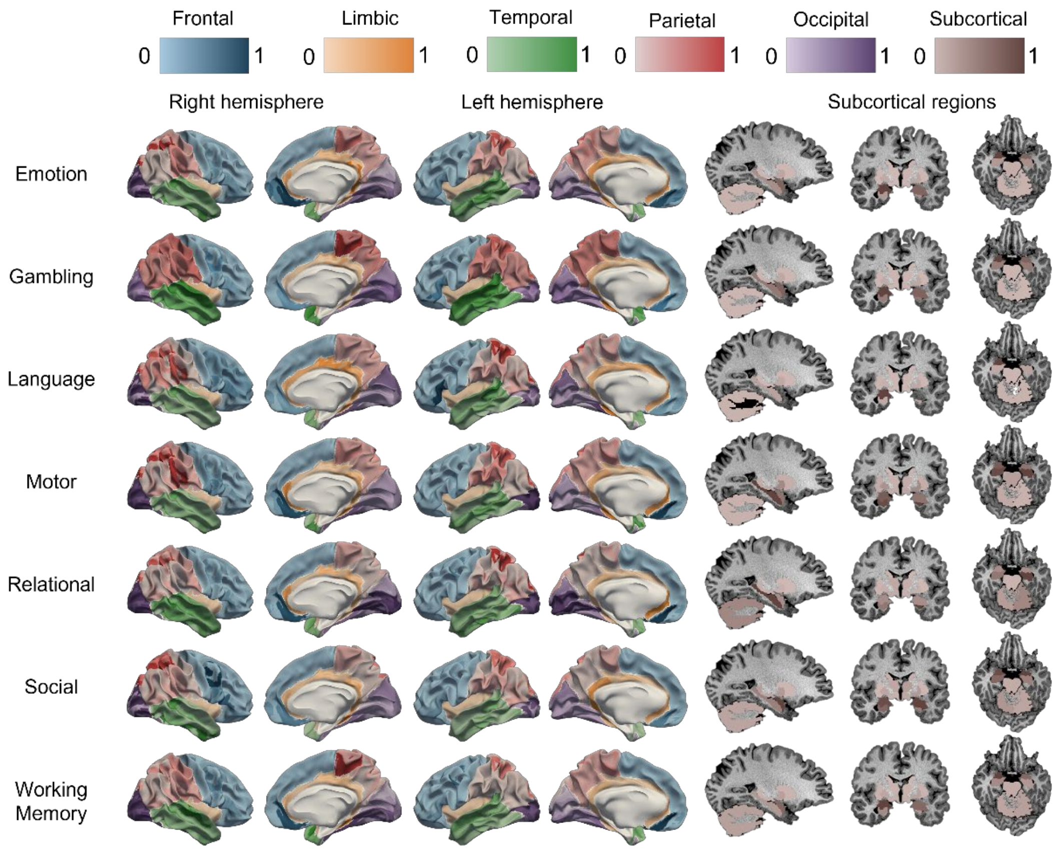


**Supplemental Fig. 2 Distribution of the functional activity weight values in the six categories based on the CHCP dataset and DK atlas.** Each category (five lobes and one subcortical area) is represented by a unique color while larger/smaller weight values correspond to darker/lighter color.

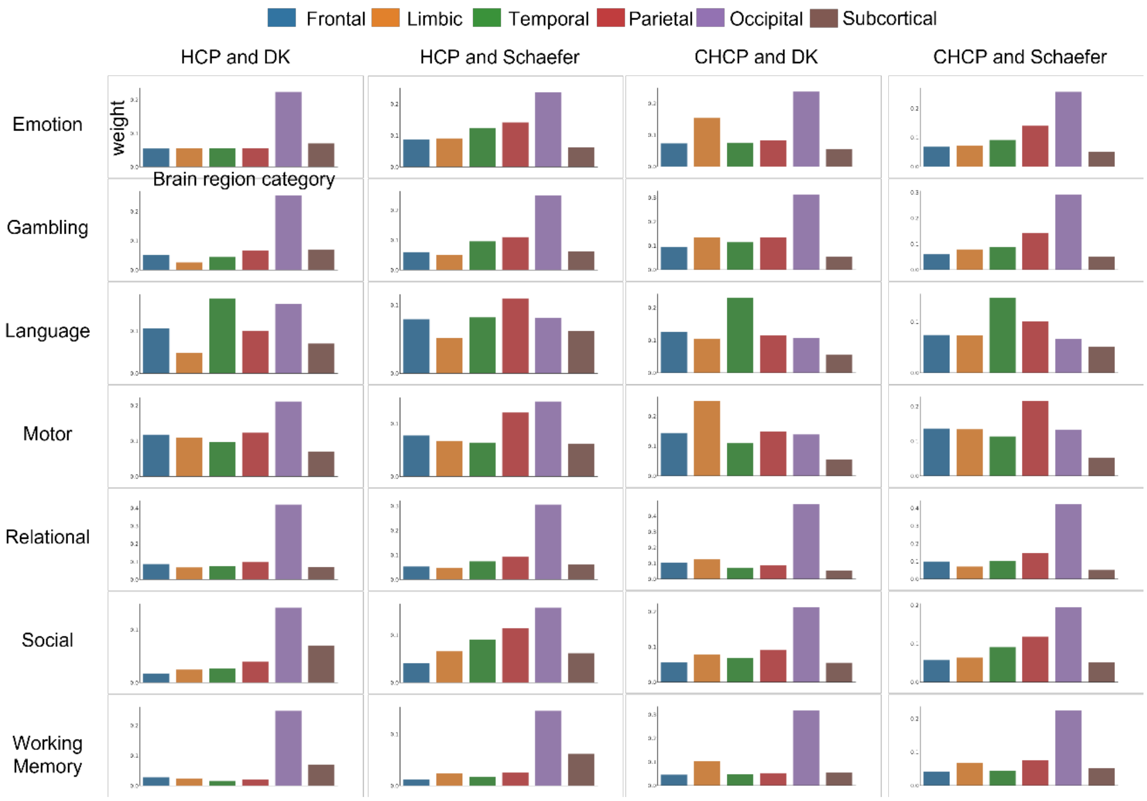


**Supplemental Fig. 3 The functional activity weight values of each category (five lobes and one subcortical area) in the identified functional neuroanatomy fingerprint based on different datasets (HCP and CHCP) and brain atlases (DK and Schaefer).** In each subfigure, the horizontal axis represents the brain region category, and the vertical axis represents the mean functional activity weight value.

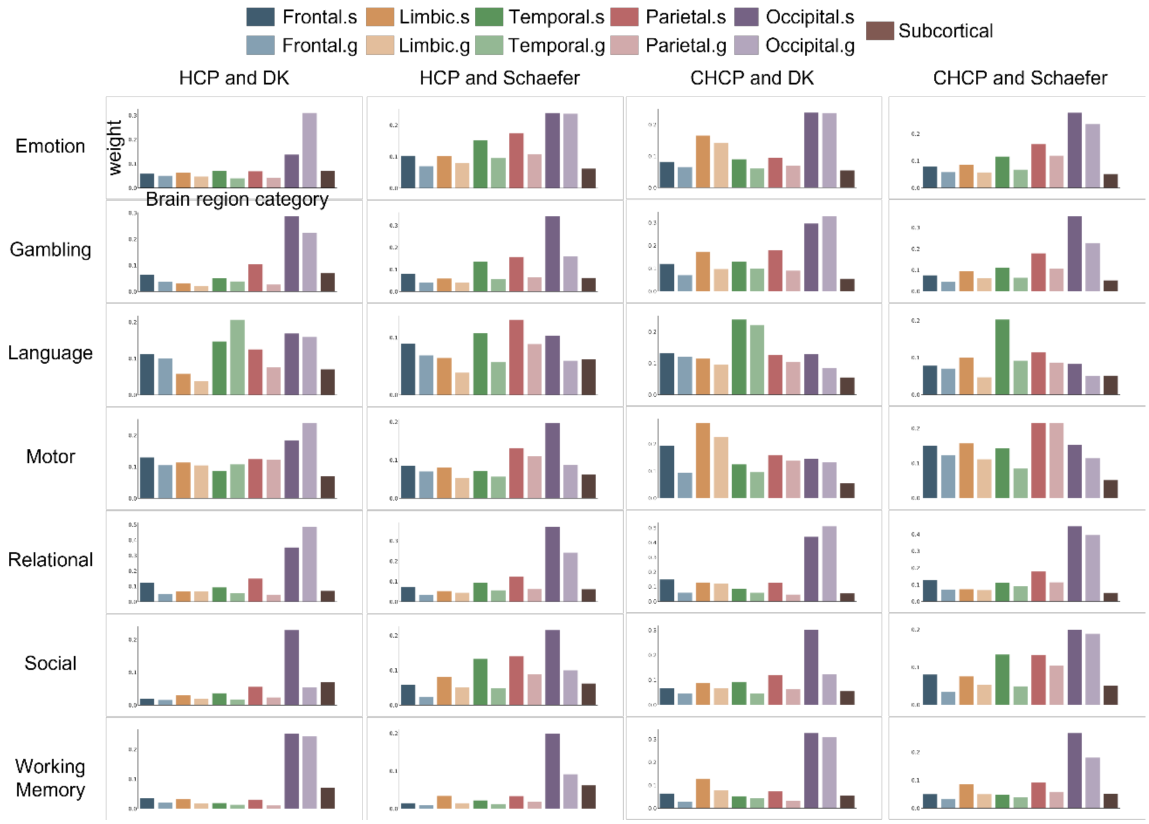


**Supplemental Fig. 4 The functional activity weight values of eleven categories (gyri/sulci part of each of the five lobes plus one subcortical area) in the identified functional neuroanatomy fingerprint based on different datasets (HCP and CHCP) and brain atlases (DK and Schaefer).** In each subfigure, the horizontal axis represents the brain region category, and the vertical axis represents the mean functional activity weight value. s and g denote sulci and gyri, respectively.

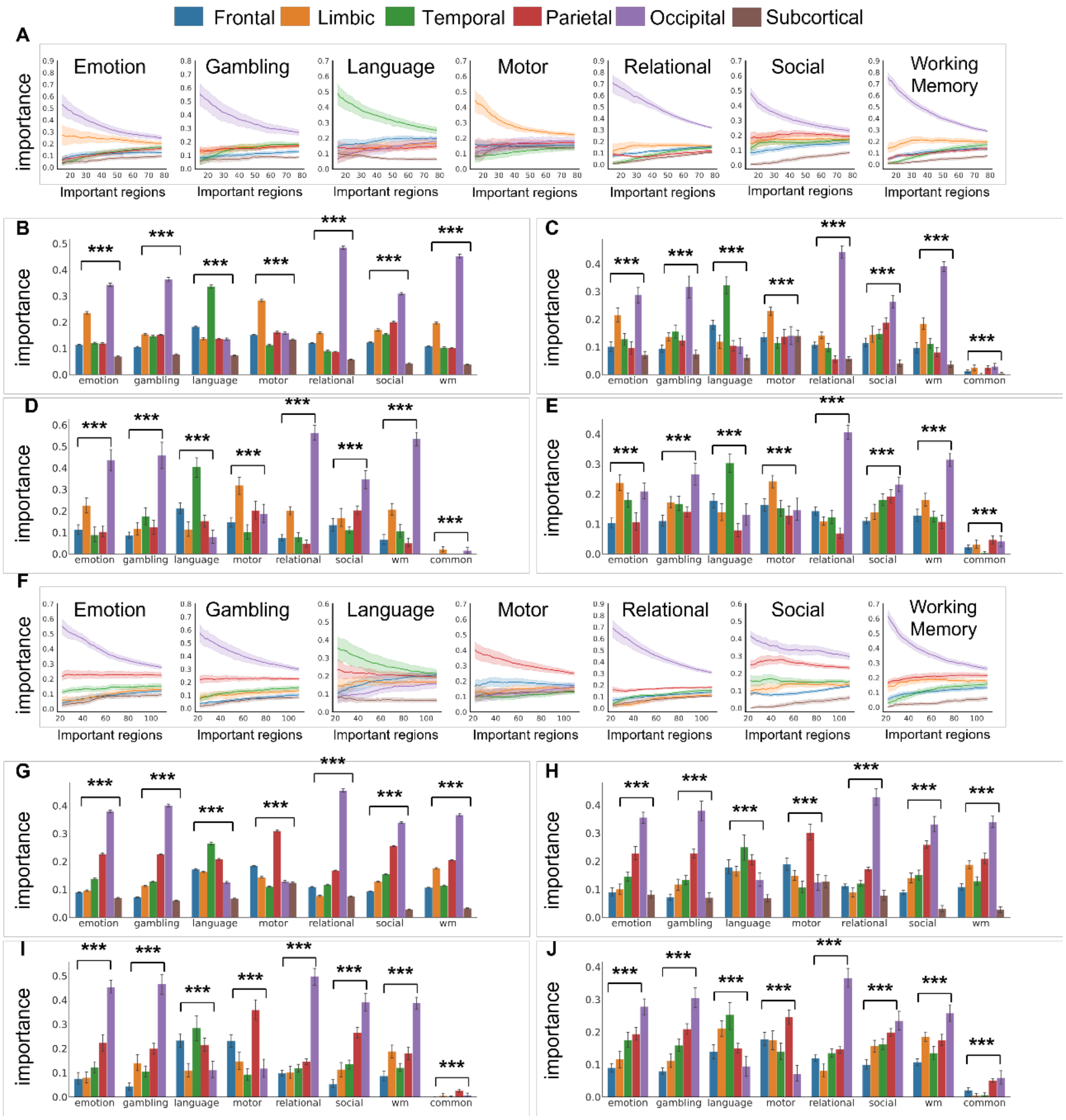


**Supplemental Fig. 5 Distribution of 'importance' among the six categories using CHCP dataset and DK/Schaefer brain atlases. (A, F)** Comparisons of 'importance' among the six categories in each mental process under different thresholds of important brain regions from top 10% (16 brain regions) to top 50% (80 brain regions) with a 0.5% interval using DK and Schaefer atlas, respectively. **(B, G)** Statistical comparison of 'importance' among the six categories in each mental process in (A, F), respectively. **(C, H)** Comparison of 'importance' among the six categories in each mental process using top 30% of important brain regions. **(D, I)** Comparison of 'importance' of gyri among the five cortical lobes in the unique and common regions, respectively using top 30% of important brain regions. **(E, J)** Comparison of 'importance' of sulci among the five cortical lobes in the unique and common regions, respectively using top 30% of important brain regions. The data is represented as the mean value with 95% confidence intervals, *** indicates $p<0.001$ in multi-way ANOVA. FDR correction is used for multiple comparisons.

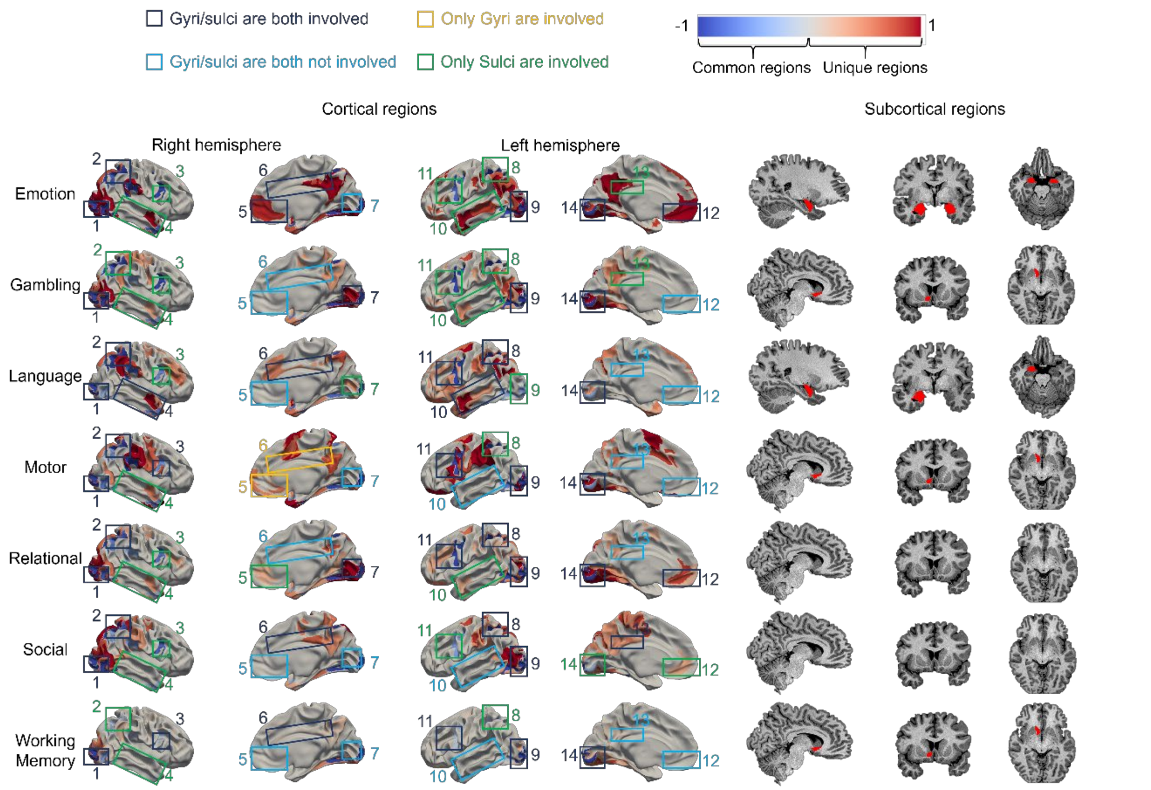


**Supplemental Fig. 6 Distribution of the common (exist in all seven mental processes) and unique (not in all seven ones) neuroanatomy regions using CHCP dataset and DK brain atlas.** The common and unique regions are colored in blue and red respectively. Top *30%* important regions with largest functional activity weight values are used as example. Fourteen representative regions in each of the four categories based on the involvement of gyri/sulci are highlighted using four different colored boxes.

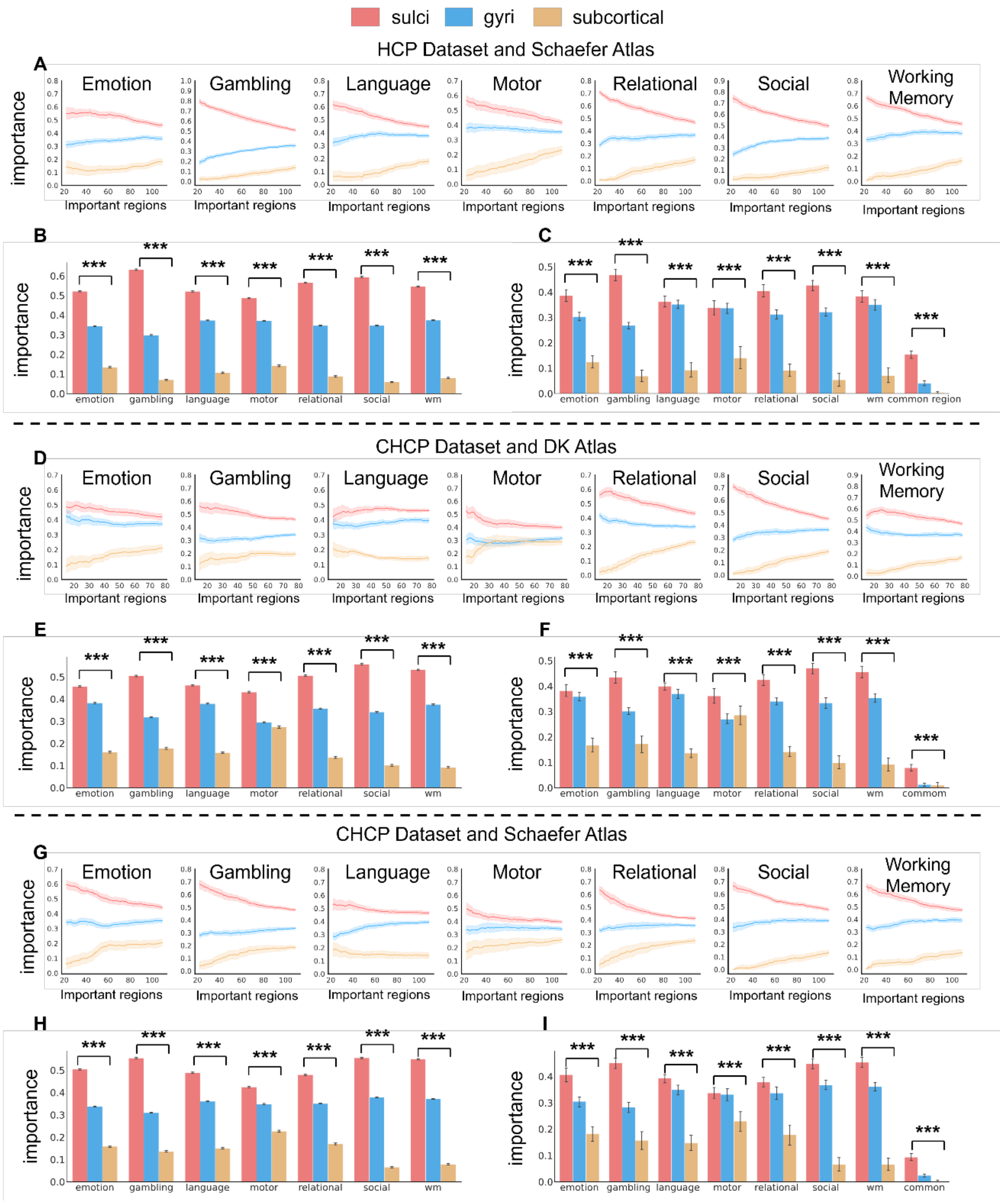


**Supplemental Fig. 7 Importance of gyri/sulci/subcortical regions in the seven mental processes. (A, D, G)** Comparison of 'importance' value among gyri, sulci, and subcortical area in each of the mental process under different thresholds of important brain regions from top 10% (16 brain regions) to top 50% (80 brain regions) with a 0.5% interval. **(B, E, H)** Statistical comparison of 'importance' among gyri, sulci, and subcortical area in each mental process in (A, D, G), respectively. **(C, F, I)** Comparison of 'importance' value among gyri, sulci, and subcortical area in each of the mental process using top 30% of important brain regions. The bars show the mean values with 95% confidence intervals, *** indicates p<0.001, and ns indicates not significant in multi-way ANOVA, respectively. The data is represented as the mean value with 95% confidence intervals, *** indicates p<0.001 in multi-way ANOVA. FDR correction is used for multiple comparisons.

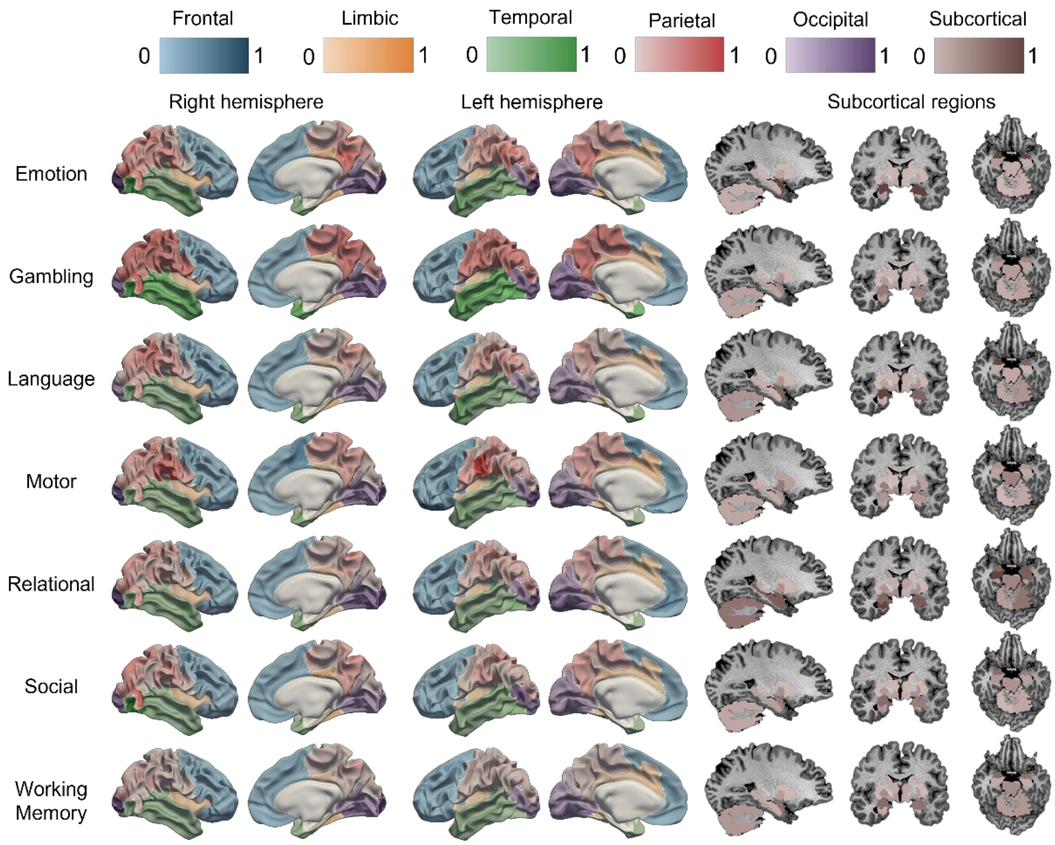


**Supplemental Fig. 8 Distribution of the functional activity weight values in the six categories based on the HCP dataset and Schaefer atlas.** Each category (five lobes and one subcortical area) is represented by a unique color while larger/smaller weight values correspond to darker/lighter color.

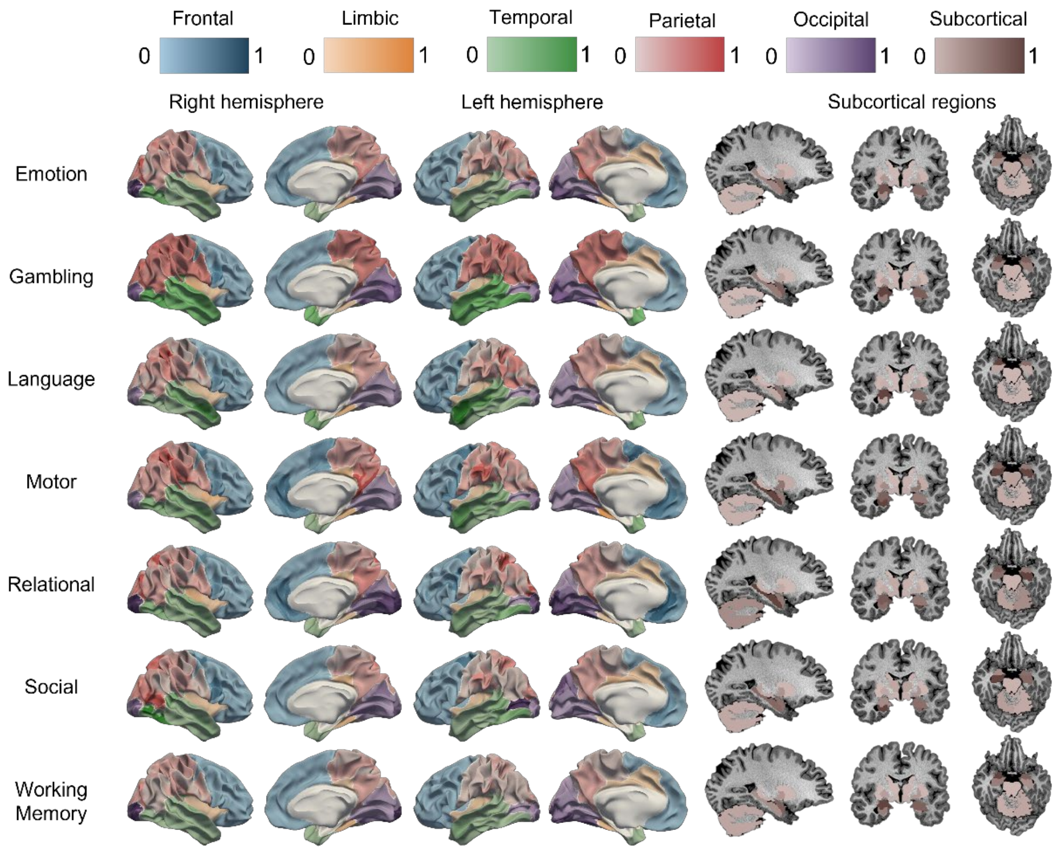


**Supplemental Fig. 9 Distribution of the functional activity weight values in the six categories based on the CHCP dataset and Schaefer atlas.** Each category (five lobes and one subcortical area) is represented by a unique color while larger/smaller weight values correspond to darker/lighter color.

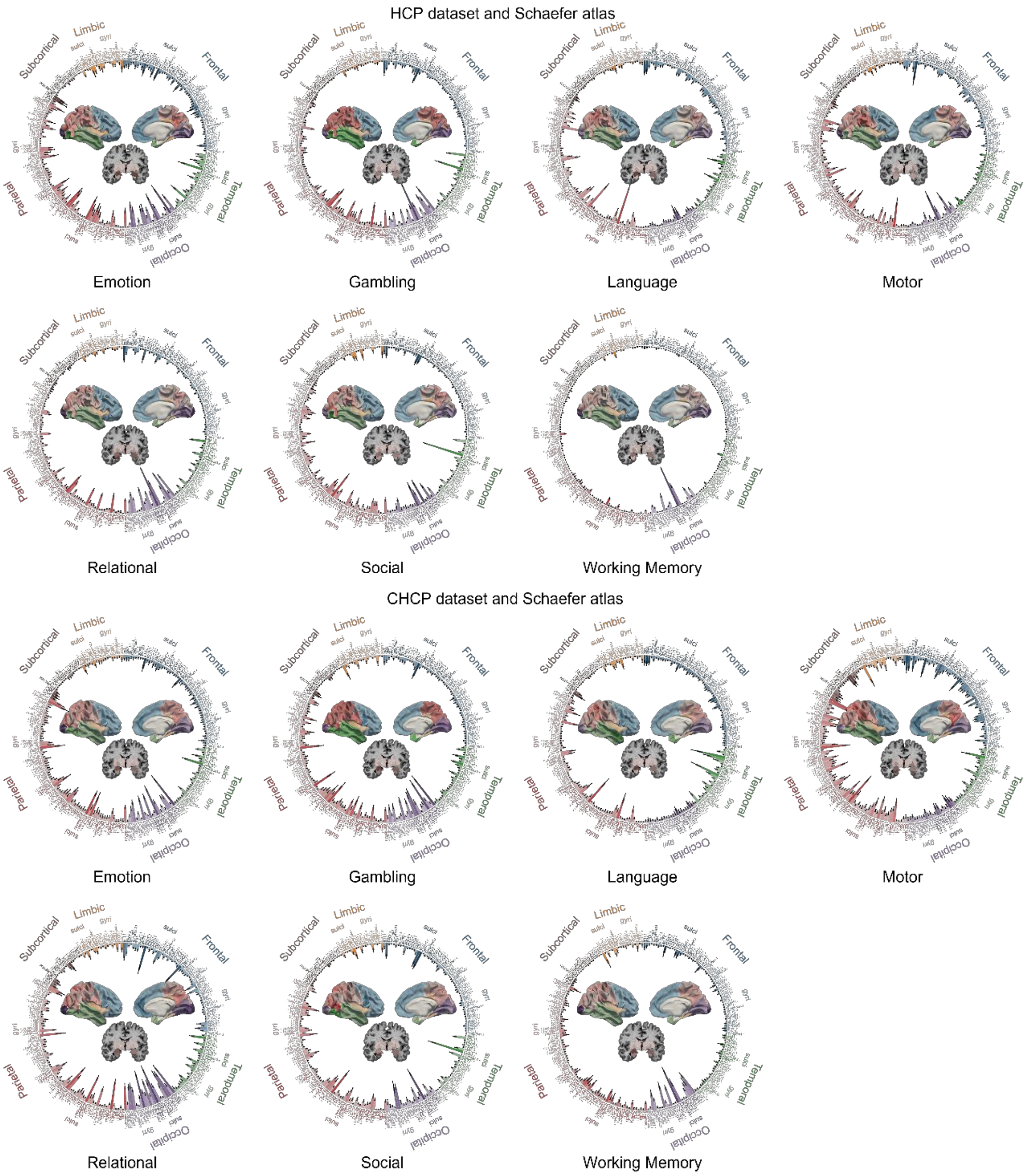


**Supplemental Fig. 10 Representation of the functional neuroanatomy fingerprint in each mental process.** The fingerprint is as a set of functional activity weight values of all neuroanatomical regions defined by the Schaefer brain atlas in HCP and CHCP dataset, respectively. Note that each value is represented as mean±std based on the thirty trained models.

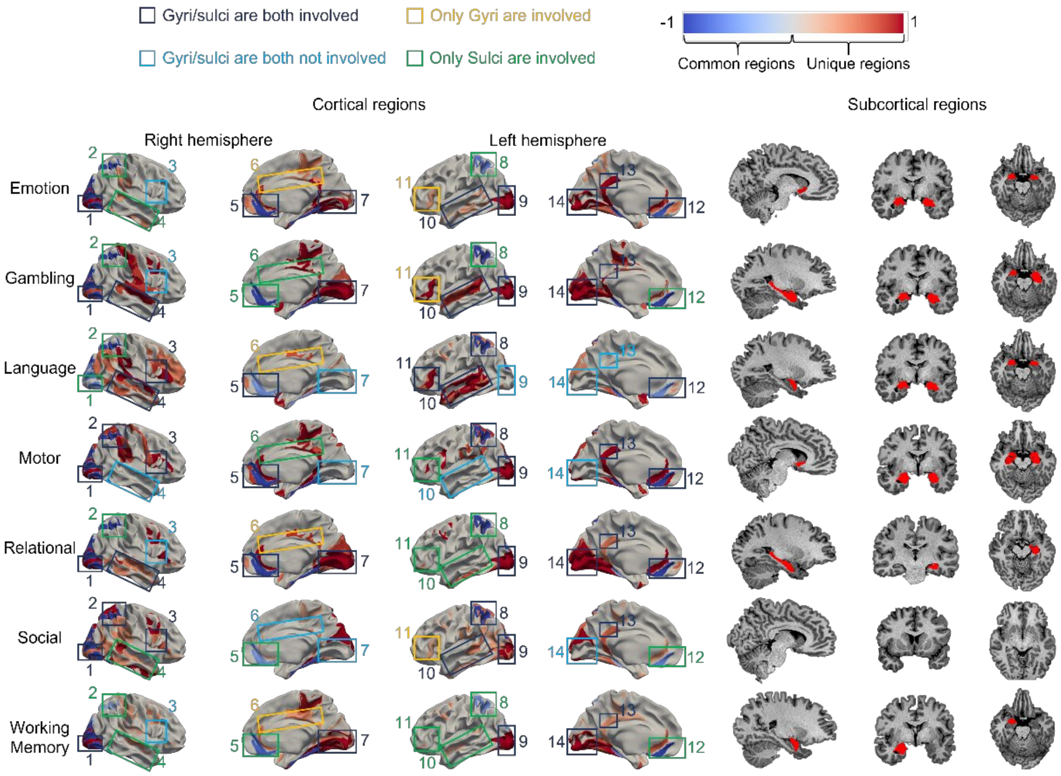


**Supplemental Fig. 11 Distribution of the common (exist in all seven mental processes) and unique (not in all seven ones) neuroanatomy regions using HCP dataset and Schaefer brain atlas.** The common and unique regions are colored in blue and red respectively. Top *30%* important regions with largest functional activity weight values are used as example. Fourteen representative regions in each of the four categories based on the involvement of gyri/sulci are highlighted using four different colored boxes.

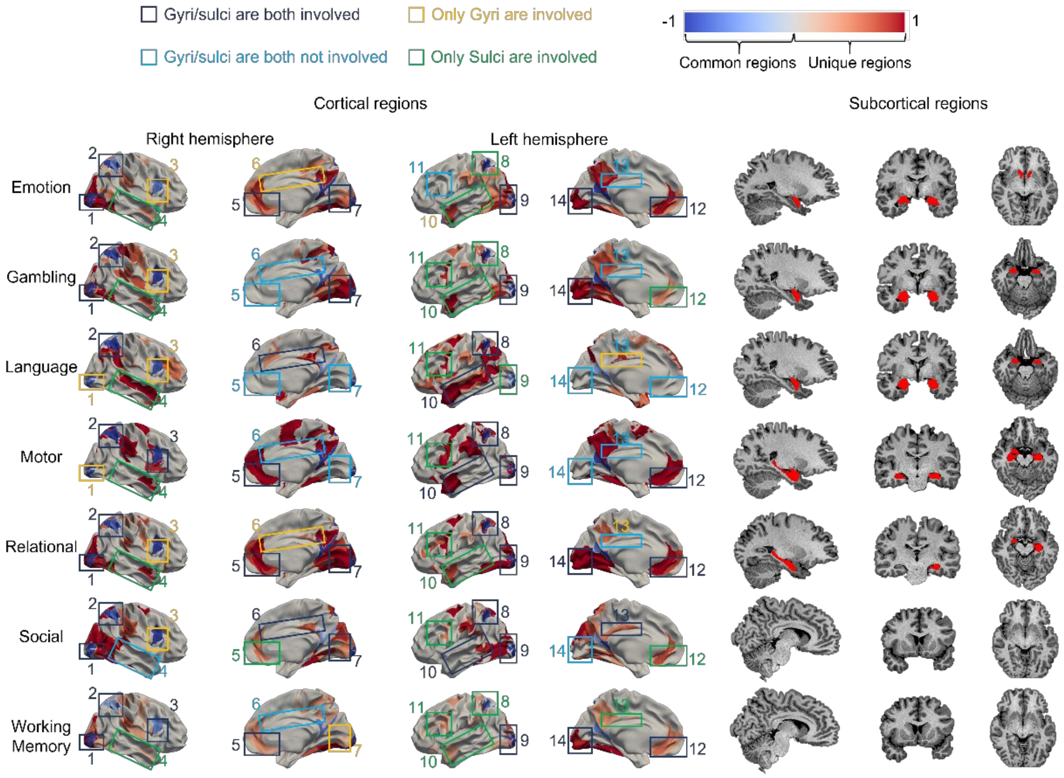


**Supplemental Fig. 12 Distribution of the common (exist in all seven mental processes) and unique (not in all seven ones) neuroanatomy regions using CHCP dataset and Schaefer brain atlas.** The common and unique regions are colored in blue and red respectively. Top *30%* important regions with largest functional activity weight values are used as example. Fourteen representative regions in each of the four categories based on the involvement of gyri/sulci are highlighted using four different colored boxes.

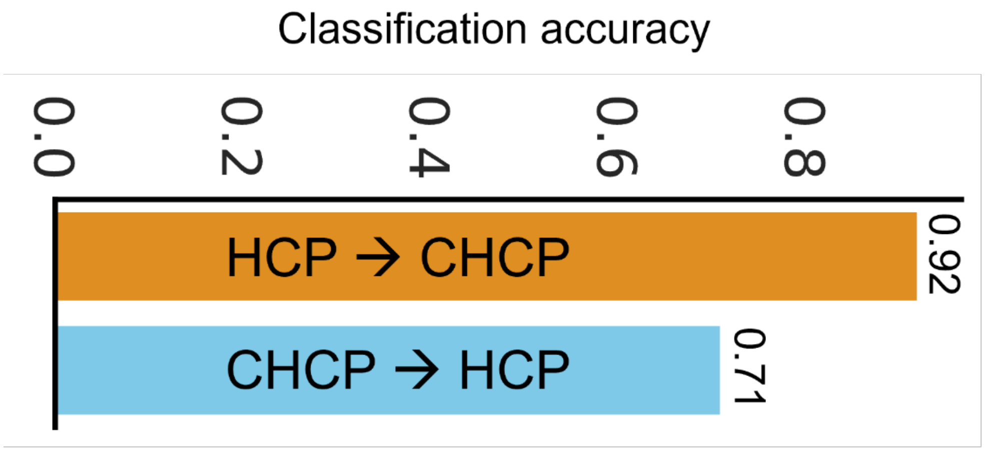


**Supplemental Fig. 13 Zero parameter update of classification between the two datasets using Schaefer brain atlas.** We directly apply the model trained on HCP dataset without any parameter updates to classify CHCP dataset, and vice versa.

**Supplemental Table 1** The multi-task classification model architecture based on DK brain atlas

| Layer | Input Shape | Connected Layer | Output Shape | Parameter |
|---|---|---|---|---|
| Input | [155, 155] | Patch embed | [1, 155, 768] | 119040 |
| Patch embed | [1,155, 768] | Position embed | [1, 156, 768] | 119808 |
| Position embed | [1, 156, 768] | Layer Norm | [1, 156, 768] | 768 |
| Norm1: Layer Norm | [1, 156, 768] | Norm1: Attention(q, k, v) | [1, 156, 2304] | 1769472 |
| Norm1: Attention(q, k, v) | [1, 156, 2304] | Norm1: Linear(q, k, v) | [1, 156, 768] | 589824 |
| Norm1: Linear(q, k, v) | [1, 156, 768] | Norm2: Layer Norm | [1, 156, 768] | 119808 |
| Norm2: Layer Norm | [1, 156, 768] | Norm2: MLP (fc1) | [1, 156, 3072] | 2359296 |
| Norm2: MLP (fc1) | [1, 156, 3072] | Norm2: MLP (fc2) | [1, 156, 768] | 2359296 |
| MLP (fc2) | [1, 156, 768] | Output | [7] | 5383 |
| One block: Norm1+Norm2<br>Total blocks: 12<br>Heads: 12 | | | | Total parameters: 85301767 |

**Supplemental Table 2** The multi-task classification model architecture based on Schaefer brain atlas

| Layer | Input Shape | Connected Layer | Output Shape | Parameter |
|---|---|---|---|---|
| Input | [219, 219] | Patch embed | [1, 219, 768] | 168192 |
| Patch embed | [1, 219, 768] | Position embed | [1, 220, 768] | 168960 |
| Position embed | [1, 220, 768] | Layer Norm | [1, 220, 768] | 768 |
| Norm1: Layer Norm | [1, 220, 768] | Norm1: Attention(q, k, v) | [1, 220, 2304] | 1769472 |
| Norm1: Attention(q, k, v) | [1, 220, 2304] | Norm1: Linear(q, k, v) | [1, 220, 768] | 589824 |
| Norm1: Linear(q, k, v) | [1, 220, 768] | Norm2: Layer Norm | [1, 220, 768] | 119808 |
| Norm2: Layer Norm | [1, 220, 768] | Norm2: MLP (fc1) | [1, 156, 3072] | 2359296 |
| Norm2: MLP (fc1) | [1, 220, 3072] | Norm2: MLP (fc2) | [1, 220, 768] | 2359296 |
| MLP (fc2) | [1, 220, 768] | Output | [7] | 5383 |
| One block: Norm1+Norm2<br>Total blocks: 12<br>Heads: 12 | | | | Total parameters: 85400071 |
